# A Spectrum of Possibilities: A Systematic Evaluation of Fluorescent Proteins in Cyanobacteria

**DOI:** 10.64898/2026.05.18.725961

**Authors:** Dennis Hasenklever, Joëlle Boecker, Angelina Grankin, Fatma Şener, Ilka M. Axmann, Anna Behle

## Abstract

Fluorescent reporters cover a wide range of applications in both basic and applied research. Whether a study involves microscopic imaging to study (co)-localization of proteins, FRET, biosensing, or quantifying gene expression, fluorophores are attractive reporter candidates due to their relatively straightforward *in vivo* readout. For microbiological applications, a wide variety of fluorescent proteins with varying excitation and emission wavelengths, brightness levels, and maturation times are available. Careful consideration is required when selecting from this large suite of proteins, especially when choosing multiple fluorophores. This is further complicated in phototrophic organisms, which exhibit strong autofluorescence, especially towards the red part of the spectrum, effectively eliminating common candidates such as mCherry. In this study, the specific properties and performance of a selection of fluorescent proteins are systematically evaluated against the background of photosynthetic pigment-derived autofluorescence in the cyanobacterium *Synechocystis* sp. PCC 6803. Specific readouts of different combinations of fluorescent proteins are also analyzed using high-throughput methods, namely plate reader fluorescent scans and single-cell flow cytometry to quantify fluorescence. The ultimate goal is to assess each fluorescent protein with regard to: 1.) Its ability to be discerned from cyanobacterial autofluorescence. 2.) Its compatibility with other fluorophores in this context. 3.) Its overall suitability in cyanobacterial research. Several highly suitable fluorescent proteins for use in cyanobacteria are identified, including mTagBFP2, mNeonGreen and mScarlet-I and suitable combinations, covering nearly the whole spectrum of visible light. This study expands the knowledge and toolset for current and future researchers and uncovers a whole spectrum of possibilities for fluorescent protein selection in cyanobacterial cell biology.

## Introduction

First isolated from the jellyfish *Aequorea victoria*, fluorescent proteins (FPs) are classified by their ability to absorb photons at a characteristic range of wavelengths, and re-emit them at a slightly higher wavelength, thereby generating a quantifiable optical output. This biophysical property (also referred to as a Stokes’ Shift) makes FPs popular reporter proteins for many *in vivo* applications in multiple cell types from diverse organisms. Since the first isolation of green fluorescent protein (GFP) in the 1960s, the repertoire of available FP colors and properties has since increased drastically, with more than a thousand FPs catalogued on FP-base [1]. Previous works involving FPs and cyanobacteria often featured green or yellow variants, which are particularly bright and thus easier to detect against cyanobacterial autofluorescence. Although cyan variants have been used in cyanobacterial research as well, their low brightness and poor signal-to-background ratios limit their application to quantitative gene expression studies. However, engineered cyan FP (CFP) variants, such as mTurquoise with improved brightness, have proven suitable for microscopy-based studies [2, 3]. Orange and red variants are rarely reported in cyanobacterial research, even though red FPs (RFPs) such as mCherry have been shown to work well in higher plants [4, 5] and green algae [6]. The reason for this is pigmentation: In addition to various chlorophylls, cyanobacteria contain phycobiliproteins (particularly phycocyanin and phycoerythrin), which exhibit strong autofluorescence with emission maximum around 575 nm (phycoerythrin), 610 nm (phycocyanin), and 660 nm (chlorophyll a) [7, 8]. Additionally, chlorophyll a can be excited between 400 and 425 nm, further reducing spectral windows available for exciting heterologous fluorophores without interference. In fact, recent studies leveraged the cyanobacterial light-harvesting complex, developing a farred FP based on an allophycocyanin subunit with a brightness level that is comparable to enhanced GFP (eGFP) [9]. Furthermore, cyanobacteriochromes have been successfully used for engineering of red/green optogenetic switches [10]. Red and farred FPs are particularly of interest in tissue cultures due to the fact that long-wavelength light exhibits more favorable penetration [11]. Cyanobacteria also possess carotenoids, with one of their function being absorption and quenching of excess energy from photosynthesis. This may also affect measurements of fluorophores in cyanobacteria, potentially reducing the effective measurable signal from a heterologously expressed FP.

FPs are commonly used to characterize genetic tools such as promoters or ribosome binding sites (RBSs) [12, 13, 14, 15]. However, dual FP systems enable more sophisticated analyses, for example, to test termination efficiencies, dual yellow FP (YFP)/ blue FP (BFP) readouts have been used to measure control (YFP) vs. readthrough (BFP) expression [16, 17]. The only reported example for orange or red FPs being used in cyanobacteria combines mScarlet-I with GFP to track two separate plasmids in *Synechococcus elongatus* PCC 7942, though only microscopy data is shown for mScarlet-I fluorescence [18]. Other studies highlight the importance of genetic context by comparing RBS strengths through readout of different FPs. While in some cases, a strong consistency was observed across different FPs [12], others have reported significant differences between FPs [19]. In a similar fashion, two different GFP variants were compared across four different cyanobacterial species using two different promoter modules, yielding drastically different results depending on the species or genetic context [20]. In one study, an oxygen-insensitive FP was required to generate oxygen biosensors, rendering standard FPs unsuitable [21]. Instead, an oxygen-insensitive, blue flavin mononucleotide–based FP (FbFP) [22] was successfully used.

Localization studies also employ FPs, particularly when investigating *in vivo* interactions between different proteins. For imaging purposes, additional factors such as photostability, maturation times, or aggregation properties need to be taken into account when selecting the optimal FP. Depending on the abundance of the protein being observed, brightness becomes a critical factor. The green FP mNeonGreen [23] has been a popular choice for these reasons [24]. One prominent study tracked *de novo* carboxysome assembly by visualizing YFP-tagged Rubisco *via* time-lapse microscopy [25]. A similar study used mTurquoise-tagged Rubisco, simultaneously monitoring mNeonGreen-tagged components of proteins involved in the spatial distribution thereof [24].

Finally, certain advanced analytical applications require at least two robust fluorescent reporters: Firstly, Foerster resonance energy transfer (FRET) enables visualization of the spatial proximity of two interaction partners through excitation of a donor FP, transfer of emission energy from the donor FP to an acceptor FP, and subsequent quenching of the donor fluorescence, thereby probing possible protein-protein interactions [26]. While an extensive database of suitable FRET pairs is available on fpbase.org [1], suitability must be determined for the individual application and target organism. The popular CFP/YFP FRET pair has been successfully applied in cyanobacterial research, including an intramolecular biosensor to study the cyanobacterial P_II_ protein [27]. Secondly, Matryoshka biosensors feature two FPs: One for sensing and one for normalizing to total protein levels. This provides a high dynamic range while circumventing biological artifacts such as expression levels [28]. Large Stokes Shift (LSS) proteins are particularly useful here, as two distinct fluorophores can be excited by the same wavelength while still emitting at different wavelengths. This also has the additional advantage of multiplexing even more FPs.

With an increasing number of studies on synthetic co-cultures on the horizon, use of FPs for monitoring individual co-culture partners also requires careful selection, as this is dependent on factors like individual growth rates, maturation times, and autofluorescence [29, 30].

In this work, 14 different FPs were systematically expressed with emission spectra ranging from 400 to 800 nm under the control of a strong, constitutive promoter in the cyanobacterium *Synechocystis* sp. PCC 6803 (*Synechocystis* hereafter). Measurements were performed in a plate reader under conditions optimized for the respective FP, as well as in a single-cell flow cytometer. Except for mCherry and mKate2, which emit near cyanobacterial autofluorescence, all FPs produced well-distinguishable signals in both assays. In the plate reader measurements, green and yellow FPs achieved the highest signal-to-background ratios_plate reader_ (ratio_PR_), particularly mNeonGreen, followed by the blue FPs, especially mTagBFP2, while in the cytometry measurements, mTagBFP2 showed the highest signal-to-background ratio_cytometer_ (ratio_cyt_), followed by the green and yellow FPs. The emission peak of LSSmOrange was clearly distinguishable from the autofluorescence, yielding a signal-to-background ratio_PR_ similar to the BFP variants in plate reader measurements. Despite spectral proximity to autofluorescence, both LSSmApple and mScarlet-I proved suitable red FP candidates. Different combinations of FPs were also assessed in terms of signal separation and spill-over. Of the candidates tested, the most suitable pairs were cyan and yellow, blue and green, or green and red/orange; however, depending on measurement settings available, blue and yellow, blue and red/orange, or yellow and red/orange may also be suitable candidates. This work aims to serve as a valuable resource for choosing the ideal FP candidate(s) based on application, equipment, and strain.

## Material & Methods

### Plasmids and Strains

All constructs tested in *Synechocystis* are based on the RSF1010 backbone pSHDY with the mobAY24F mutation [31]. A base plasmid carrying P_cpc560_ [32] upstream and the T1-T7 double terminator downstream of the NeoBrick sites was constructed. The coding sequence (CDS) of each FP was amplified with primers carrying overhangs for the strong synthetic RBS* [33] and the NeoBrick prefix upstream, as well as the NeoBrick suffix downstream of the coding sequence, and inserted using NeoBrick-specific restriction enzymes [31] and T4-Ligase. In addition, a KanR cassette was inserted in reverse orientation. The final base construct served as empty vector control (EVC) throughout the study. For dual reporter constructs, the same PCR fragments were inserted into backbones upstream or downstream of the other FP CDS in the NeoBrick suffix, resulting in dual-FP operons with the same RBS* upstream of each CDS. All constructs used in this work are summarized in Table 1. In some cases, FP CDS were codon-optimized for *Synechocystis* sp. PCC 6803 (mVenus, mCerulean, mTagBFP2) using the IDT codon-optimization tool, or manually for *Synechococcus elongatus* PCC 7942 (mNeonGreen, [34]). All DNA and amino acid sequences used in this work can be found in Supplementary File S3 and S4, respectively. For *Synechocystis*, plasmids were mobilized *via* triparental mating using *Escherichia coli* strain NEB5*α* carrying the RSF1010 cargo plasmid, J53 carrying the RP4 plasmid required for conjugative transfer, and a *Synechocystis* sp. PCC 6803 non-motile, glucose-tolerant wild type from Uppsala, Sweden as the recipient strain [35].

**Table 1:**
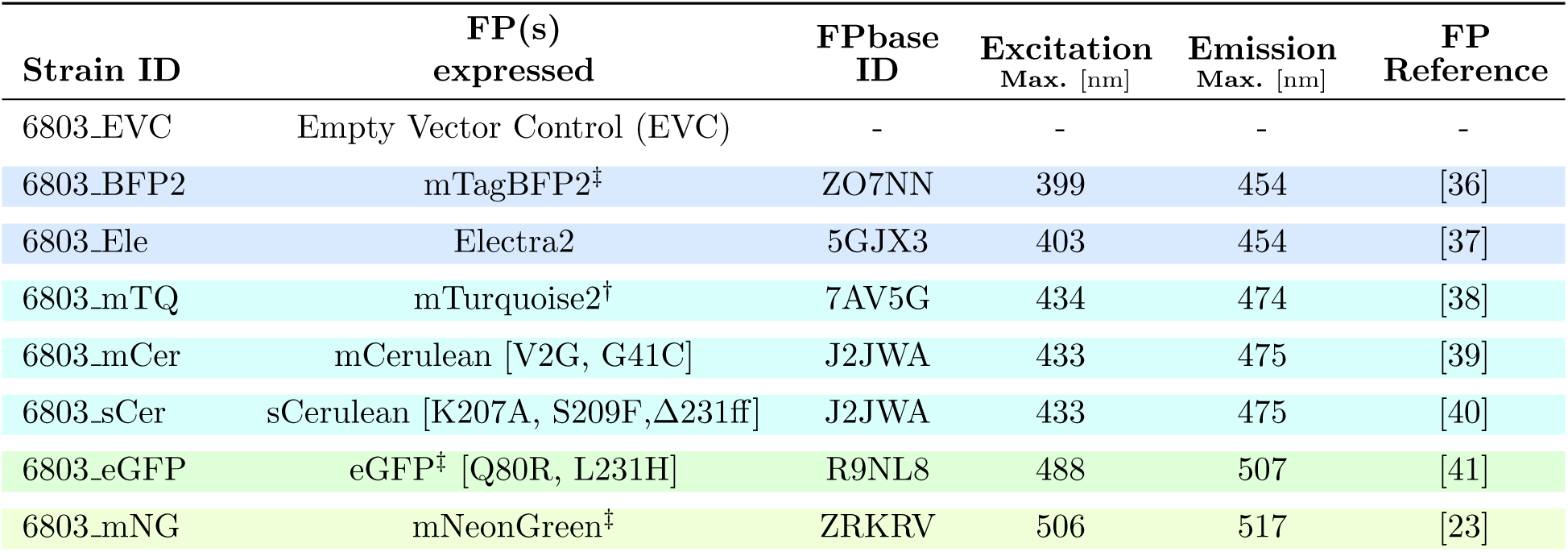
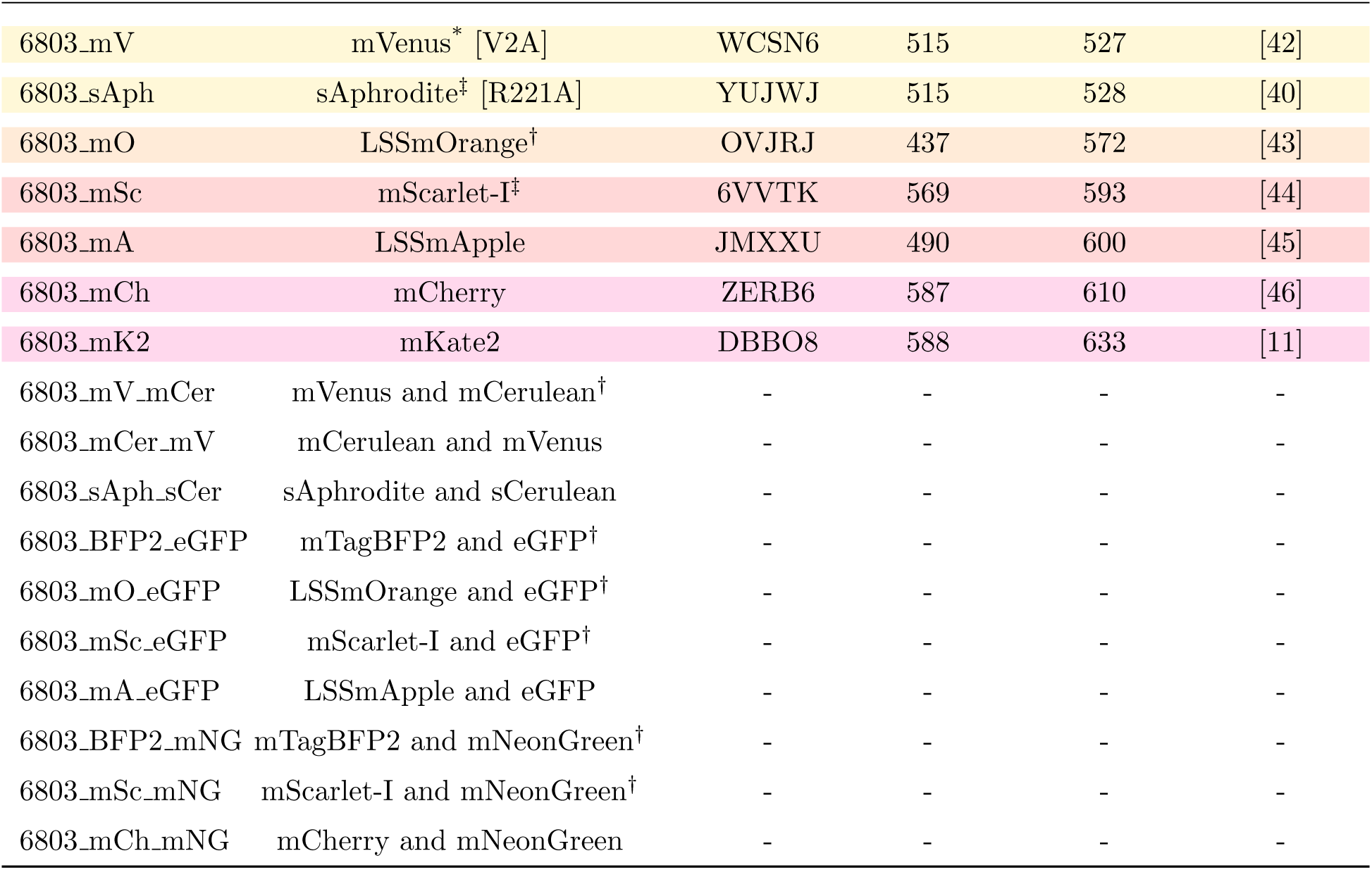
FP constructs tested in this work. EVC: Empty vector control. Amino acid exchanges are listed in brackets, if applicable, in relation to the official amino acid sequence entry found on FPbase. Δ231ff denotes a truncation from amino acid 231 to the end of the ORF, in reference to the respective FPbase ID. All strains were constructed in this study. ^*^most suitable for plate reader, ^†^most suitable for cytometry, ^‡^ suitable for both. Suitability was determined per color.

### Bacterial cultivation conditions

BG11 medium was modified after [47] using 0.521 mg/L (1.4 µM) Na_2_EDTA · 2H_2_O instead of 1 mg/L (2.8 µM) MgNa_2_EDTA and 30 mg/L (131 µM) instead of 40 mg/L (175 µM) K_2_HPO_4_ · 3H_2_O. Additionally, 10 mM TES buffer from a 1 M stock solution adjusted to pH 8 at 30 °C was used. In the final medium, the pH was around pH 7.6 at 30 °C and atmospheric conditions. All components except for CuSO_4_ and Fe-ammonium-citrate were added before autoclaving. All cyanobacterial strains generated in this work were separately restreaked on fresh BG11 plates containing Spectinomycin (20 µg · mL*^−^*^1^) from cryopreserved samples. These antibiotic concentrations were supplemented throughout all experiments. Plates were maintained in an Infors Multitron Pro incubator at 30 °C supplemented with 35 µmol photons · m*^−^*^2^·s*^−^*^1^ white light illumination (14 % setting), 0.5 % CO_2_ and 50 % humidity for 7 - 10 days. For liquid cultivation, each 5 mL of 1x BG11 was inoculated in 6-well plates (TPP 92006, Merck) and shaken in a Labwit Ultimate Cell USOK 1257 incubator at 150 rpm (25 mm throw) and 30 °C, supplemented with 80 µmol photons · m*^−^*^2^·s*^−^*^1^ white light illumination (36 % setting), 0.5 % CO_2_ and 50 % humidity. When precultures turned turbid (3 days), each culture was backdiluted to an optical density at 750 nm (OD_750_) of 0.2 and transferred to 12-well plates in quadruplicates of 1 mL. Plates were placed on a shaker with a microplate attachment (KS 15A, Edmund Buehler, Germany, 17 mm throw), which was operated at 200 rpm, alternating between 1 hour clockwise and 5 minutes counterclockwise. Cell cultures were grown for an additional two days in Infors Multitron Pro incubator at 30 °C supplemented with 35 µmol photons · m*^−^*^2^·s*^−^*^1^ white light illumination (14 % setting), 0.5 % CO_2_ supply and 75 % humidity. Light intensities were calibrated using a LI-COR Quantum Q 111763 sensor directly on the sticky plate of the incubator. The final OD_750_ of the cultures was around 3 and comparable for all strains. Samples were diluted in fresh BG11 medium to an OD_750_ of 1.0 in 220 µL and transferred to a black 96-well plate with a clear base (4titude, UK, 4ti-0221) for plate reader measurements. From this, 20 µL were diluted 10-fold in fresh BG11 medium in a clear 96-well plate for single-cell flow cytometry measurements (final volume of 200 µL for each plate).

### Microplate Reader settings and measurements

Samples of 200 µL were loaded into a black-walled, clear-bottom 96-well plate. Measurements were performed in a BMG CLARIOstar plate reader (SN: 430-1281). Excitation and emission scans of all samples were performed in 5 nm increments with 10 nm bandwith around excitation and emission wavelength and separate excitation wavelength and gain settings for each strain. In addition, an absorbance scan including the absorbance at 750 nm was recorded for all wells. A detailed description of the specific settings regarding excitation and emission wavelengths, as well as gain settings, can be found in the Supplementary Methods, Supplementary File S1.

### Flow cytometry settings and measurements

Measurements were performed with a Beckman Coulter CytoFLEX S Flow Cytometer (Model No.: B75442, SN: BE51180) using the auto acquisition plate mode and clear 96-well plates with 5 seconds backflush and 5 seconds mixing between each sample. To cover as many different settings as possible, three different detector configurations were chosen. A detailed description of all laser and bandwidth filter settings and the three detector configurations can be found in Supplementary File S2. Briefly, the letters describe the excitation laser (V=Violet: 405 nm; B=Blue: 488 nm; Y=Yellow: 561 nm; R=Red: 638 nm), -H and -A stand for height and area of the signal peak, respectively, while numbers describe the emission wavelength and bandwidth of the filters (e.g. R660-10-A means: red laser, 655 - 665 nm bandwidth, area). All gains were set to 500 for fluorescence channels and 25 for the forward scatter (FSC) and side scatter (SSC). The threshold for detection was set to 1,500 in the forward scatter height (FSC-H) channel. After gating for cyanobacterial red autofluorescence in the histogram of the R660-10-H (or APC-H) channel using a vertical divider gate *>*10^4^ arbitrary units [A.U.], 50,000 cyanobacterial cells were recorded per sample with a sample flow rate of 10 µL min^-1^, around 2,000 events per second and an abort rate of around 2 %. For each detector configuration, one of the culture replicates in a separate well was measured.

## Data Analysis

### Microplate Reader measurements

All fluorescence measurements were normalized to the absorbance at 750 nm (OD_750_) from the absorbance spectra. Emission spectra of each FP and corresponding EVC within the same dataset were feature-scaled using its minimum and maximum fluorescence intensities (Eq.1). Signal-to-background ratios_PR_ were calculated according to Eq. 2. The maximum ratio was determined for each FP (Eq. 3),

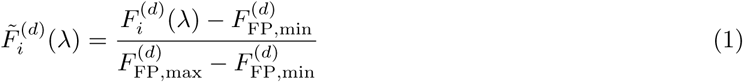

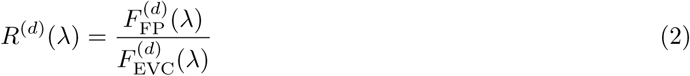

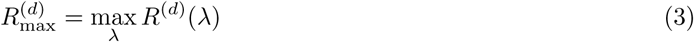

where 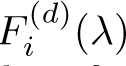 denotes the fluorescence intensity of strain *i* at wavelength *λ* for dataset *d*, and 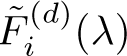 the corresponding feature-scaled fluorescence obtained by min–max normalization based on the FP spectrum, with *i* ∈ {FP, EVC}.

### Flow cytometry

For data analysis of the cytometry data, gating was performed in two steps. First, gating for cyanobacterial red autofluorescence in the histogram of the R660-10-H (or APC-H) channel was performed using a vertical divider gate *>*10^4^ A.U., leaving the recorded 50,000 cyanobacterial cells. Second, singlets (single cells) were gated by their size in the histogram of the FSC-Width channel using a vertical divider gate *<*10^3^ A.U.. These two gates were combined to cyanobacterial singlet cells leaving roughly 49,500 events per sample. All calculations were performed for the cyanobacterial singlet cells and the area values (-A) of the fluorescence channels. The median and percentiles needed for the violin plot were calculated for all gated cyanobacterial singlet cells from each strain population, individually. These median fluorescent intensities (MFI) were used to calculate ratios of the FP-strain population divided by the respective EVC population within the same fluorescence channel (MFI_FP_/MFI_EVC_), and referred to as signal-to-background ratio_cyt_ in the text.

### Data processing

Plate Reader data was analyzed using Python 3.12.10. Cytometry data was analyzed using the FlowCytometryTools package [48] in Python 3.9.18. All python code was deposited in the same Annotated Research Context (ARC) as the raw data and will be made available once peer-reviewed.

## Results & Discussion

### Choice of FPs and construct design

A range of FPs with varying excitation and emission wavelengths was selected, covering the entire spectrum of visible light (Figure S1). In order to gain a higher experimental resolution in the blue and red range, multiple candidates were chosen, often (partially) overlapping in their biophysical properties. Two large stokes’ shift candidates, LSSmOrange and LSSmApple, were also included. All candidates and their biophysical properties, as well as their origin, are summarized in Table 1. *Synechocystis* strains were designed to express each FP from an RSF1010-based, self-replicating plasmid and the strong constitutive P_cpc560_ promoter. Strain construction is detailed in the Materials & Methods section, all constructs and deviations in their amino acid sequence are summarized in Table 1.

### The choice of fluorophores is restricted in cyanobacteria due to their prominent autofluorescence

For each FP, an optimal excitation wavelength was selected, based on the following criteria: (1) Published values, (2) inclusion of peak emission fluorescence in spectrum, (3) minimal distance between excitation and emission spectra. After optimizing individual gains for each strain (based either on the excited fluorophore or the highest overall fluorescence overall), a spectral emission scan was performed of all strains in each separate setting to identify the optimal emission wavelength with minimal distance between the excitation and start of the emission scan. Figure S1 shows the excitation and emission spectra for each fluorophore at their optimal excitation wavelength, as well as representative cyanobacterial pigments allophycocyanin (APC) and chlorophyll a.

To assess autofluorescence levels when exciting the different FPs, an excitation scan of the wild type (WT) *Synechocystis* strain was performed (Figure S2). Two different emission wavelengths were chosen based on the phycocyanin and chlorophyll a emission maxima (Figure S2 A, B, respectively). While ultraviolet (UV) and blue ranges showed minimal signal, autofluorescence increased substantially at excitation wavelengths above 500 nm. None of the red FPs investigated in this work are excited at the maximum excitation wavelength, but a significant co-excitation is to be expected nonetheless. To further highlight this, emission scans for two representative excitation wavelengths were also performed (Figure S2 C, D), which show the same trend towards high background fluorescence at 600 nm and above.

Due to high autofluorescence signals of the BG11 blank, especially in the blue and cyan range, blank-corrected values of the EVC were often negative, making the calculation and visualization of signal-to-background ratio_PR_ and emission spectra difficult (Figure S4). The raw, OD-corrected fluorescence of FP strains vs. the EVC was therefore compared in the following chapters.

### Blue and cyan FPs

In total, five FP variants were selected from the spectral range between 400 and 435 nm for excitation. Among them, two BFP variants (excitation around 400 nm) were tested: mTagBFP2 (previously shown to perform adequately in cyanobacteria [49]) and Electra2 (engineered for enhanced brightness [37]) Moreover, three distinct cyan variants (excitation around 435 nm) were included in the experimental setup: mCerulean, sCerulean and mTurquoise2. Both BFPs are well-distinguishable from the non-FP-expressing control in whole-cell emission scans (Figure 1 A, B). Electra2, which was reported to be even brighter than mTagBFP2 in some animal models [37], shows a weaker signal compared to mTagBFP2. This is also reflected in the signal-to-background ratio_PR_. In contrast, mCerulean, sCerulean and mTurquoise2 showed nearly identical emission spectra and signal-to-background ratios_PR_ (Figure 1 C, D; Figure S3 A). While CFPs are notoriously dim due to their low quantum yield, mTurquoise2 was developed specifically towards a high quantum yield and improved brightness *in vitro*. However, other factors, such as expression level, protein stability, or intracellular conditions may contribute to intracellular brightness *in vivo*. Overall, both BFP variants outperform the CFPs in terms of being distinguishable from the EVC.

**Figure 1:**
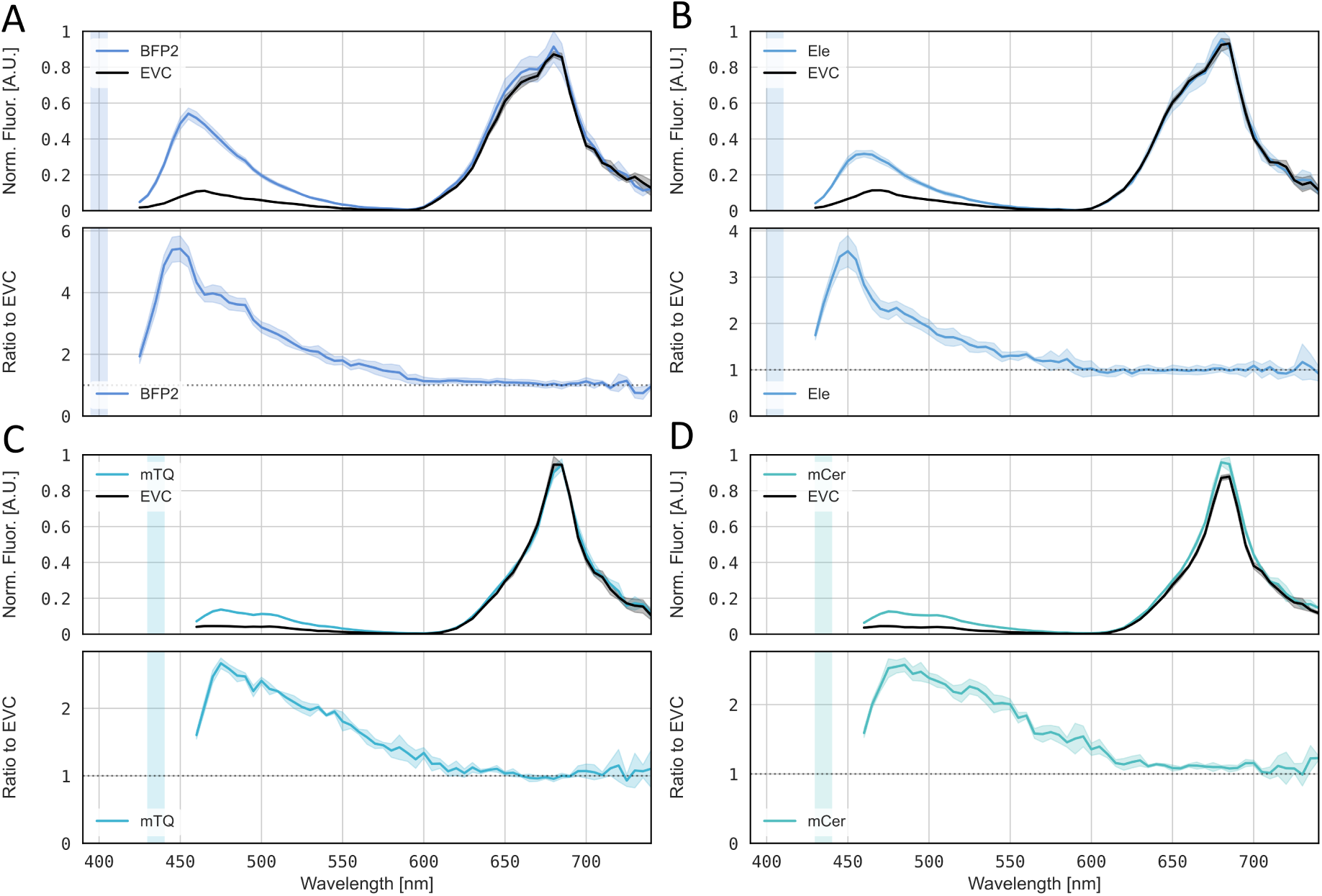
Whole-cell emission scans of blue and cyan single-FP strains tested in this work. Norm. Fluor. [A.U.]: Normalized Fluorescence [arbitrary units]. A: BFP2 (mTagBFP2), B: Ele (Electra2), C: mTQ (mTurquoise2), D: mCer (mCerulean). A,B,C,D top panels: Whole-cell emission scan of *Synechocystis* expressing FP of interest. For each condition, the empty vector control (EVC) strain was measured alongside. Raw data were normalized to OD_750_. Data for each plot (FP + respective EVC in black) was scaled ranging from 0 to 1. The shaded region shows the standard deviation of four analyzed replicates. A,B,C,D bottom panels: Signal-to-background ratio_PR_ of FP-encoding strain to EVC; the shaded region shows the standard deviation of four analyzed replicates. The optimal range of excitation wavelengths used for each FP according to published data is marked by a vertical shaded bar.

### Green and yellow FPs

Green and yellow FPs were represented by two selected variants each. While eGFP is among the earliest modifications of the original *Aequorea victoria* GFP (avGFP) [50], the variant used in this work has two additional mutations [41]. mNeonGreen was developed more recently from a different FP origin (LanYFP), and is reported to be one of the brightest green FPs [23]. This FP was included because of its popularity in various cyanobacterial expression and protein localization studies [51, 34]. mVenus was selected for similar reasons. In addition, sAphrodite, a codon-modified version of Venus with the same excitation and emission spectra [40], was included.

In contrast to the blue and cyan FPs, the green and yellow FPs enable an extremely high dynamic range, both through low cyanobacterial autofluorescence around their emission maxima and their high signal intensities (Figure S2; Figure 2). Accordingly, the green FPs emit at higher signal levels than the autofluorescence in the 600 - 750 nm range. Both yellow FP variants showed strong signal-to-background ratio_PR_, though sAphrodite exhibited a higher maximum signal, likely due to improved expression levels. These high signal-to-background ratios_PR_ were achieved despite excitation outside the maximum (Supplementary Methods, Figure S1). Due to the close proximity of excitation and emission spectra and maxima of green and yellow FPs, great care should be taken if selecting one of these for analysis in the plate reader. This combination overall accounts for the high signal-to-background ratio_PR_ observed for all green and yellow variants, certainly contributing to the explanation why these are most commonly used in cyanobacteria.

**Figure 2:**
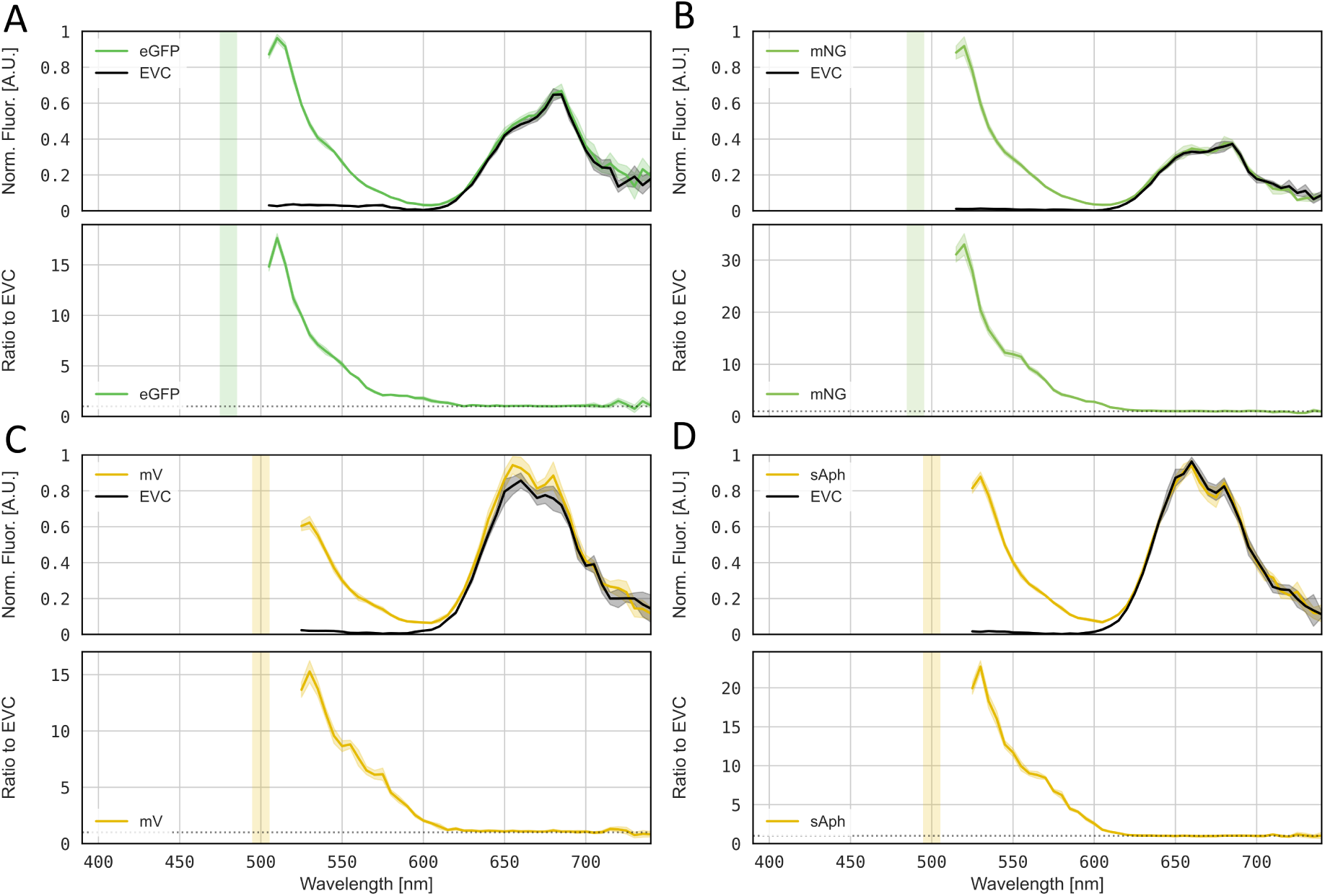
Whole cell emission scans of green and yellow single-FP strains tested in this work. Norm. Fluor. [A.U.]: Normalized Fluorescence [arbitrary units]. A: eGFP, B: mNG (mNeonGreen), C: mV (mVenus), D: sAph (sAphrodite). A,B,C,D top panels: Whole cell emission scan of *Synechocystis* expressing FP of interest. For each condition, the empty vector control (EVC) strain was measured alongside. Raw data were normalized to OD_750_. Data for each plot (FP + respective EVC in black) was scaled ranging from 0 to 1, The shaded region shows the standard deviation of four analyzed replicates. A,B,C,D bottom panels: Signal-to-background ratio_PR_ of FP-encoding strain to EVC; the shaded region shows the standard deviation of four analyzed replicates. The optimal range of excitation wavelengths used for each FP according to published data is marked by a vertical shaded bar.

### Red and LSS FPs

As mentioned earlier, the use of orange and red FPs in cyanobacteria is prone to high background fluorescence due to the proximity of their maxima to the phycobiliprotein and chlorophyll autofluorescence (Figure S2 A, B, respectively). However, LSS-FPs may overcome this limitation because they are excited at wavelengths much further away from the excitation range of autofluorescent components. Indeed, both LSSmOrange and LSSmApple show a distinguishable signal in their maximum range (Figure 3 A, C), the latter partially merging into the autofluorescence peak. Both LSS-FP variants showed peak signal-to-background ratios_PR_ at their corresponding emission maxima of 575 nm or 600 nm (Figure 4). LSSmOrange, the brighter variant, also shows an overall higher ratio, making it similarly suitable as the BFP variants, whereas LSSmApple resembled CFP performance. Another likely reason for the good dynamic range of these FPs is the fact that autofluorescence is less intense when exciting at 435 or 490 nm (Figure S2). Both LSS variants tested in this work suffer from a long maturation time (over 100 min), which may need to be carefully considered depending on the application.

**Figure 3:**
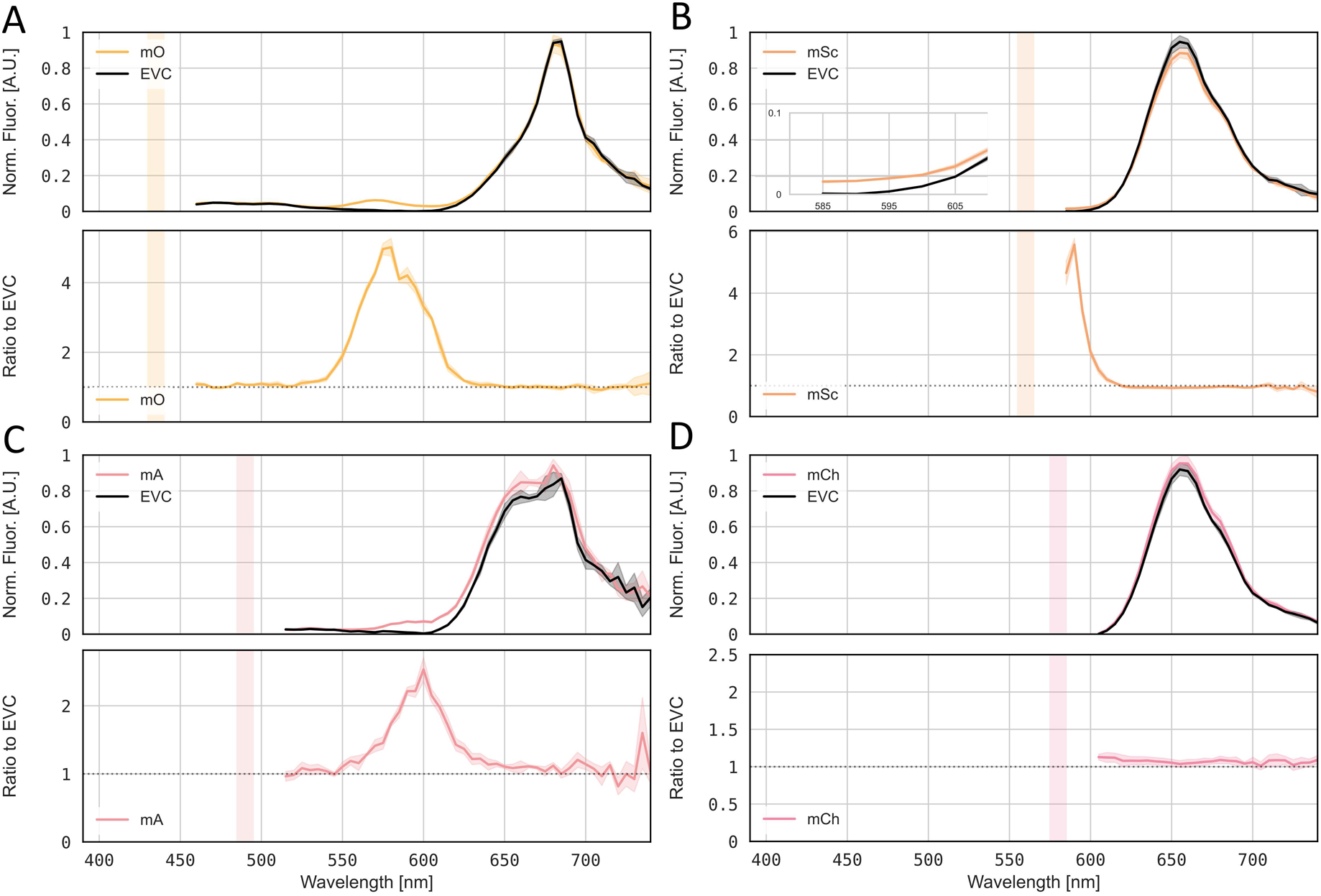
Whole-cell emission scans of LSS and red single-FP strains tested in this work. Norm. Fluor. [A.U.]: Normalized Fluorescence [arbitrary units]. A: mO (LSSmOrange), B: mSc (mScarlet-I), inset shows close-up within the relevant emission range, C: mA (LSSmApple), D: mCh (mCherry). A,B,C,D top panels: Whole-cell emission scan of *Synechocystis* expressing FP of interest. For each condition, the empty vector control (EVC) strain was measured alongside. Raw data were normalized to OD_750_. Data for each plot (FP + respective EVC in black) was scaled ranging from 0 to 1, The shaded region shows the standard deviation of four analyzed replicates. A,B,C,D bottom panels: Signal-to-background ratio_PR_ of FP-encoding strain to EVC; the shaded region shows the standard deviation of four analyzed replicates. The optimal range of excitation wavelengths used for each FP according to published data is marked by a vertical shaded bar.

**Figure 4:**
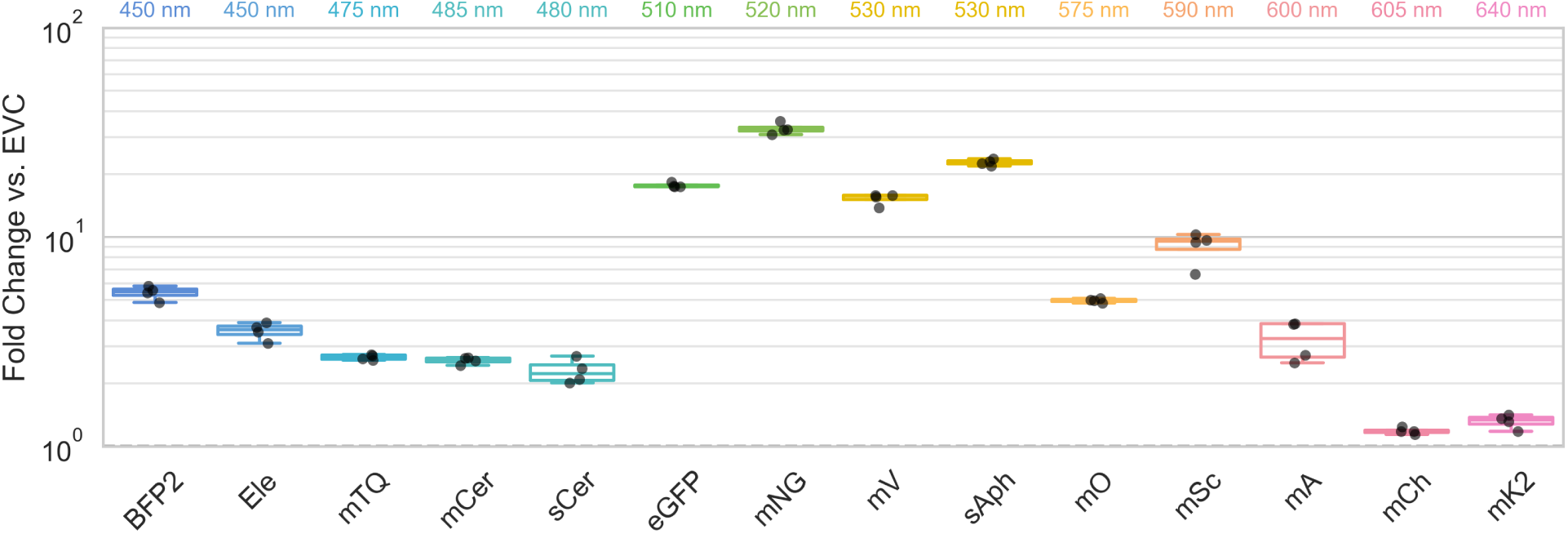
Optimal signal-to-background ratios_plate reader_ of each FP strain. The maximum signal-to-background ratio_PR_ and corresponding emission wavelength, (shown on top, +/- 5 nm) was determined for each flu-orophore at its optimal excitation/emission settings. Data represents four biological replicates. EVC: Empty vector control. BFP2: mTagBFP2. Ele: Electra2. mTQ: mTurquoise2. mCer: mCerulean. sCer: sCerulean. mNG: mNeon-Green. mV: mVenus. sAph: sAphrodite. mO: LSSmOrange. mSc: mScarlet-I. mA: LSSmApple. mCh: mCherry. mK2: mKate2.

Among the red FPs, only the mScarlet-I strain shows distinguishable fluorescence above the EVC, positioned just below the start of the autofluorescence peak (Figure 3 B). Despite the proximity, the signal-to-background ratio_PR_ of mScarlet-I at 590 nm is higher than all FPs except the yellow and green variants. On the other hand, mCherry and mKate2 emission appears to either overlap too much with the autofluorescence, or the FPs are too dim to distinguish them from the background (Figure 3 D, Figure S3 B). To summarize the findings from all emission scans, the maximum signal-to-background-ratios_PR_ were extracted for each FP and the corresponding emission wavelength (+/- 5 nm) was determined (Figure 4), revealing a clear trend towards favorable measurement conditions for yellow and green FP variants. Other suitable FPs for *Synechocystis* are mScarlet followed by LSSmOrange, the BFPs, the CFPs and LSSmApple, while RFPs in the further red part of the spectrum are unsuitable.

### Flow Cytometry enables monitoring of multiple FPs simultaneously

While limited in its flexibility in terms of FP-specific detector settings, single-cell flow cytometry is especially suitable for monitoring multiple parameters in the same population. Since individual cells are measured separately, it also eliminates biological artifacts related to culture density, a common issue in plate-reader-based measurements in cyanobacteria. It also enables the observation of heterogeneity such as differences in levels of FP expression, in the entire population. Finally, for some applications such as the high-throughput detection of individual fluorescence-tagged cell populations in co-cultures, flow cytometry is crucial and cannot easily be replaced by other methods. In contrast to the settings used for the plate reader measurements, the excitation wavelengths of the flow cytometer rely on a number of distinct lasers, in this case the four lasers violet (405 nm), blue (488 nm), yellow (561 nm) and red (638 nm), which all lie within the range of excitation of the FPs investigated in this work, but not necessarily at their optimum (Figure S1).

To assess the applicability of individual FP variants across diverse cytometry detector configurations (which vary between laboratories), all suitable and available bandpass filter-laser combinations capable of distinguishing FP strains from the EVC were tested (Supplementary Methods, Supplementary File 2). In order to compare their suitability for detection of the individual FPs, signal-to-background ratios_cyt_ were calculated by calculating median fluorescence across all cyanobacterial singlet cells within one strain and dividing those FP-strain median values by EVC median values, for all fluorescence channels across all strains. Signal-to-background ratios_cyt_ were visualized as a heat map, highlighting the best suitable channel for detection, identified by peak ratios in each column (Figure 5).

**Figure 5:**
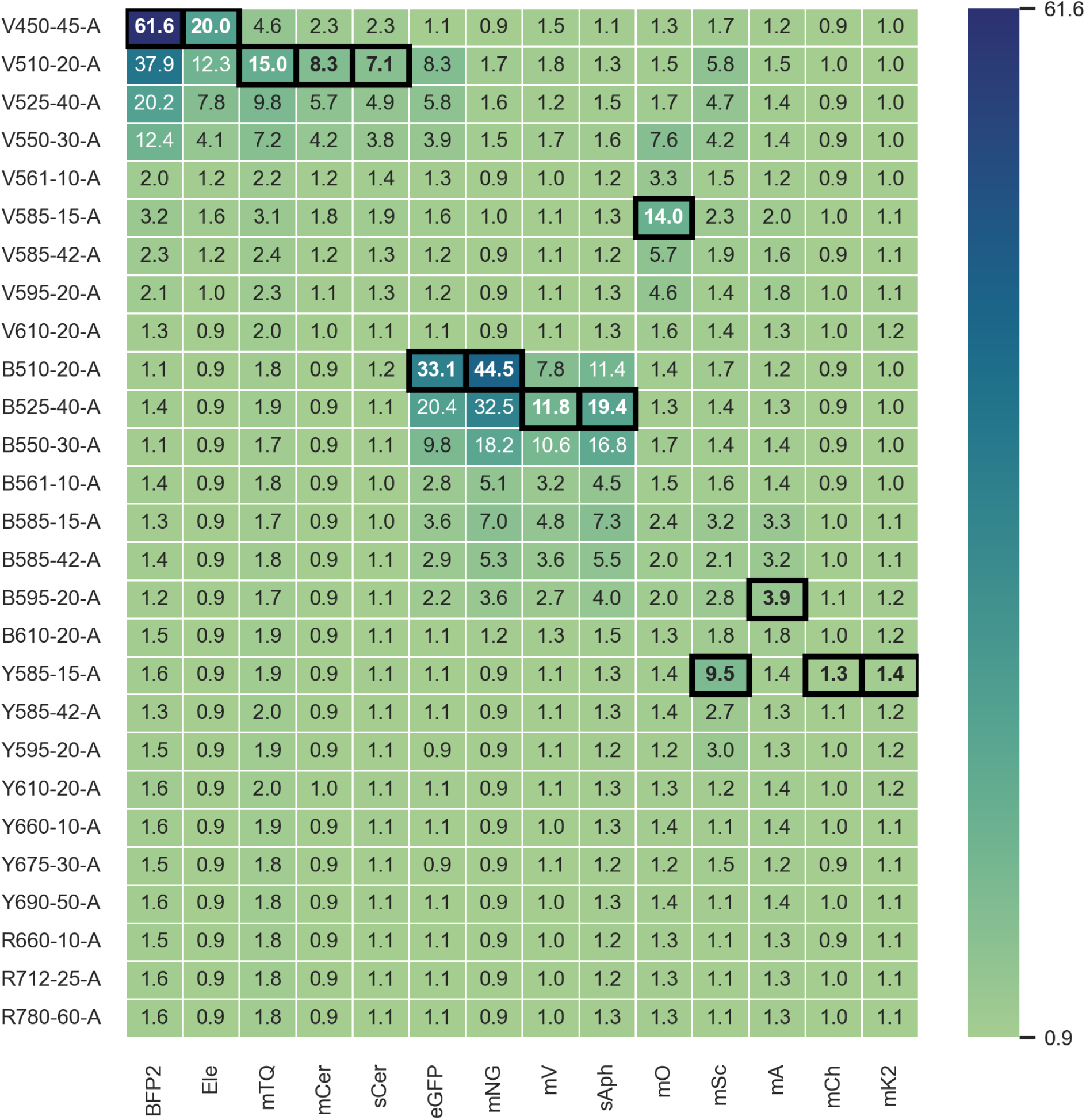
Signal-to-background ratio_cytometer_ for single FPs. The ratio of median FP/EVC (empty vec-tor control) fluorescence (signal-to-background ratio_cyt_) was determined for all combinations of single FP-strains (x-axis) and available fluorescence channels (y-axis). In the channel names, the letters describe the excitation laser (V=Violet: 405 nm; B=Blue: 488 nm; Y=Yellow: 561 nm; R=Red: 638 nm), -A stand for area of the signal peak, while numbers describe the emission wavelength and bandwidth of the filters. Color scale represents a 0.9 - 61.6 scale of the signal-to-background ratio_cyt_. Highest values determined for each separate FP strain are highlighted in bold letters and shown in black boxes. BFP2: mTagBFP2. Ele: Electra2. mTQ: mTurquoise2. mCer: mCerulean. sCer: sCerulean. mNG: mNeonGreen. mV: mVenus. sAph: sAphrodite. mO: LSSmOrange. mSc: mScarlet-I. mA: LSSmApple. mCh: mCherry. mK2: mKate2.

The blue FPs are most easily detectable in the V450-45-A channel with signal-to-background ratios_cyt_ of 61.6 for mTagBFP2 and 20.0 for Electra2, but can also be detected well in the V510-20-A, V525-40-A, and V550-30-A channels with decreasing intensity. These results are in line with the results from the fluorescence scans where the emission scans revealed a maximum emission around 454 nm and a higher signal-to-background ratios_cyt_ for mTagBFP2 compared to Electra2.

The cyan FPs are most easily detectable in the V510-20-A channel with signal-to-background ratios_cyt_ of 15.0 for mTurquoise2, 8.3 for mCerulean and 7.1 for sCerulean and can also be detected well in the V525-40-A, and V550-30-A channels with decreasing intensity, in line with what can be observed in the emission scans (Figure 1 C, D).

Surprisingly, the strongest blue and cyan FPs, mTagBFP and mTurquoise2, showed increased signal-to-background ratios_cyt_ up to 1.6 and 2.0 in the “red autofluorescence filters” (above 585 nm) in the yellow and red laser, which are not detectable with this intensity in any of the other FP strains except for mScarlet-I. Even the red FPs mCherry and mKate2, which should be visible in those filters after excitation with the yellow laser, do not reach those signal-to-background ratios_cyt_.

The green FPs are most easily detectable in the B510-20-A channel with signal-to-background ratios_cyt_ of 44.5 for mNeonGreen and 33.1 for eGFP and can also be detected well in the B525-40-A and B550-30-A channels with decreasing intensity. Similarly, the yellow FPs are most easily detectable in the B525-40-A channel with signal-to-background ratios_cyt_ of 19.4 for sAphrodite and 11.8 for mVenus and can also be detected well in the B550-30-A and B510-20-A channels with decreasing intensity. Both green and yellow FPs show increased signal-to-background ratios_cyt_ above 1 in nearly all tested channels in the blue laser up to B595-20-A although their emission maxima are in the range from 507 nm and 527 nm (Figure S1). Due to this overlap of excitation and emission, green and yellow FPs can not be reliably distinguished using the cytometer and should not be combined in dual-FP experiments.

Surprisingly, eGFP also showed increased signal-to-background ratios_cyt_ up to 8.3, which is comparable with mCerulean and even higher than sCerulean, after excitation with the violet laser (405 nm) and detection in the V510-20-A channel. As the filter V510-20-A covers the emission maximum of the green FPs, the most likely explanation is a weak excitation of eGFP at 405 nm (17 % of maximum, Figure S1) and subsequent detection at its emission maximum. On the other hand, mNeonGreen with an excitation efficiency of only 1 % at 405 nm, shows a low signal-to-background ratio_cyt_ of only 1.7 in this channel (Figure S1). So similar to YFP/GFP combinations, CFP/GFP combinations show spectral cross-talk as GFP fluorescence might be picked up in the V510-20-A channel, though the use of a green FP with a higher emission maximum, such as mNeonGreen and excitation laser separation mitigate this effect.

The red and orange FP variants are most easily detectable in the 585-15, 585-42 and 595-20 filters in the respective lasers. As for the LSS-FPs, LSSmOrange is more easily detectable with a signal-to-background ratio_cyt_ of 14.0 in the V585-15-A channel compared to LSSmApple with a signal-to-background ratio_cyt_ of only 3.9 in the B595-20-A channel. Notably, both LSS-FPs were detectable in the red filters of the respective other laser with a lower signal-to-background ratio_cyt_ around 2, enabling detection flexibility across laser lines.

Among the other red FPs, mScarlet-I is the most easily detectable FP with a signal-to-background ratio_cyt_ of 9.5 in the Y585-15-A channel. The other two RFPs, mKate2 and mCherry, are not really distinguishable from the red autofluorescence with signal-to-background ratios_cyt_ of 1.4 and 1.3 in the Y585-15-A channel, respectively. All filters in the yellow and red laser above 610-20 basically only pick up the red autofluorescence with signal-to-background ratios_cyt_ around 1, except for mTagBFP2 and mTurquoise2.

Surprisingly, mScarlet-I also shows signal-to-background ratios_cyt_ between 5.8 and 4.2 in the V510-20-A, V525-40-A, and V550-30-A channels of the violet laser, although it only has 1 % of maximal excitation (Figure S1) at 405 nm and does not really show much emission in this region of the spectrum.

Overall, the signal-to-background ratios_cyt_ are highest for the blue and green FPs followed by the yellow and cyan FPs and finally the orange and red FPs. Generally, flow cytometry yielded higher signal-to-background ratios (Figure 5) than plate reader assays (Figure 4) and also showed a slightly different order. As mentioned previously, data obtained from plate reader emission scans was not blank-corrected due to high autofluorescence of BG11, combined with potential cell-density-related quenching effects that result in negative blank-corrected values, especially in the blue range. Blank-corrected, and therefore negative or close-to-zero values (Figure S4) would have resulted in negative or artificially high fold-changes. These effects are not present in cytometry data, as single cells are individually measured and no blank-correction needs to be performed. They also may explain the discrepancies in the blue and cyan signal-to-background ratios between cytometry and plate reader data, which are much more aligned in the green, yellow, orange and red FPs, although they certainly need to be taken into account when comparing the two methods.

Based on these highest signal-to-background ratios_cyt_ optimal detection channels for each FP were selected (Figure 5). Violin plots were generated to visualize the fluorescence intensity distributions across cell populations compared to the EVC, showing a clearly identifiable peak for each FP strain with a slightly right skewed but almost symmetrical distribution on a logarithmic scale (Figure 6). Only the FP strains containing mTurquoise2 and mScarlet-I exhibit a slightly stretched peak in their distribution, which could be caused by a slight difference in expression across the population. Nonetheless, all FP-strains show only one homogeneous population, which indicates a homogeneous expression of the FP and a good stability of the genetic construct without a huge metabolic burden (which could otherwise result in escape mutations). In terms of absolute fluorescence intensity, the green, yellow and blue FP strains show the highest intensity, followed by the mScarlet- and LSSmOrange-, the cyan FP-, and the remaining red FP strains in these settings. However, these absolute intensities are not directly comparable and are insufficient to comment on the FP brightness itself, as they may also depend on molecular conditions such as the codon-usage, determining the expression level and FP content of the cell, as well as technical settings such as the gain, laser intensity, and bandpass filter bandwidth, which might also not be optimal for each FP. Decisive for the visibility of the FP is not only the absolute fluorescent intensity, but also the absolute intensity of the autofluorescence of the cell, highlighted by the EVC. This autofluorescence intensity is dependent on the wavelength of the excitation laser, as well as the wavelength of the emission being picked up by the bandpass filter. Therefore, excitation wavelengths upwards of 525 nm in the excitation spectrum, as well as emissions in the emission spectrum exceeding 600 nm, are problematic for visibility (Figure S2).

**Figure 6:**
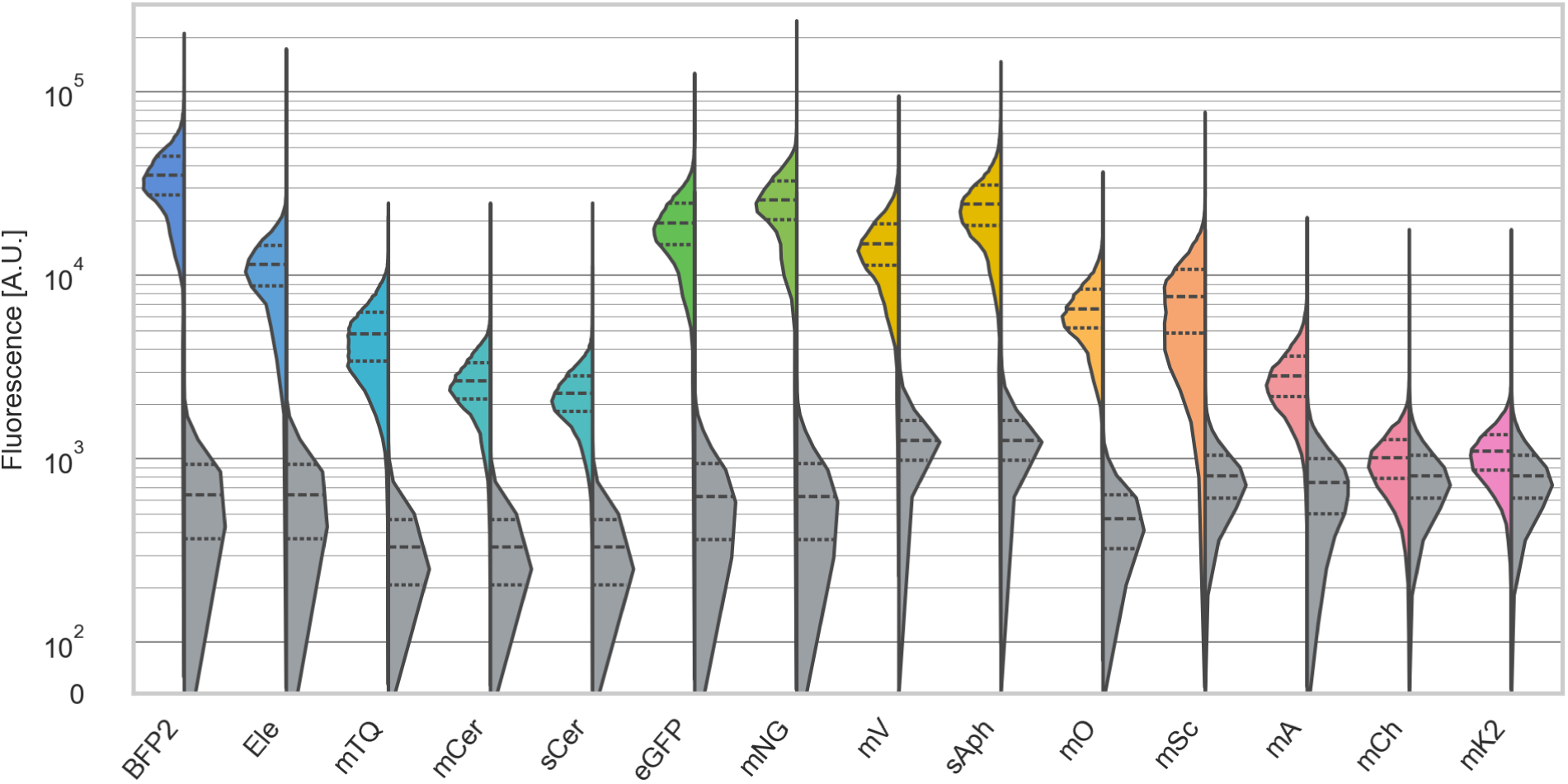
Split violin plots of single FP population compared to the EVC in its corresponding optimal channel. Violin plots show the kernel density estimation (KDE) and quartiles calculated on all cyanobacterial singlet cells (max. 50,000) for each FP population (left side of violin) and the corresponding empty vector control (EVC, right side of violin). Dashed lines represent the median (50th percentile) and dotted lines represent the 25th and 75th percentile. Violin plots are displayed on a symlog scale with a linear region from 0 - 10^2^ and a logarithmic region from 10^2^ - 3×10^5^ showing the fluorescence [arbitrary units]. In the channel names, the letters describe the excitation laser (V=Violet: 405 nm; B=Blue: 488 nm; Y=Yellow: 561 nm), -A stand for area of the signal peak, while numbers describe the emission wavelength and bandwidth of the filters. BFP2: mTagBFP2. Ele: Electra2. mTQ: mTurquoise2. mCer: mCerulean. sCer: sCerulean. mNG: mNeonGreen. mV: mVenus. sAph: sAphrodite. mO: LSSmOrange. mSc: mScarlet-I. mA: LSSmApple. mCh: mCherry. mK2: mKate2. The optimal channels for each FP are: BFP2/Ele: V450-45-A, mTQ/mCer/sCer: V510-20-A, eGFP/mNG: B510-20-A, mV/sAph: B525-40-A, mO: V585-15-A, mA: B595-20-A, mSc/mCh/mK2: Y585-15-A.

This is exactly the reason why most of the red FPs are hardly distinguishable from the red autofluorescence across nearly all excitation wavelengths, but especially in the orange and red excitation spectrum (Figure S2). For the two red FPs mCherry and mKate2, a clear differentiation or even separation from the EVC-fluorescence in a channel above the Y585-15-A channel is nearly impossible (Figure S7). Another example is the yellow FP sAphrodite, which has a similar distribution and median absolute fluorescence as mNeonGreen, but the autofluorescence in the B525-40-A channel is higher compared to the B510-20-A channel, leading to a lower overall signal-to-background ratio_cyt_ (Figure 5, Figure 6). Therefore, it is not surprising that the optimal channels with the highest signal-to-background ratio_cyt_ for all FPs exhibit comparably low autofluorescence in the EVC, with a maximal intensity of 2,000 A.U. in the B525-40-A channel. Still, except for the red FPs, these channels are also the ones covering the optimal area of the emission peak of the respective fluorophore (Figure S1).

Most of the distributions in the violin plot show a clear separation of the FP-strain from the EVC with nearly no overlap. This is particularly important for co-culture applications, where distinct cell populations are labeled with different FPs [30]. For this purpose, a clear separation using a threshold gate is beneficial. However, variants like mScarlet, LSSmApple, and also the tested cyan FPs show substantial overlap between FP and EVC populations, making them less suitable for this type of study. While the other tested red FPs mCherry and mKate2 proved fully unsuitable, the tested green, yellow and blue FPs are ideal options for dual-reporter assays, providing excellent distinction.

### A subset of dual FP combinations can be clearly distinguished from one another

Dual FP pairs were chosen and constructed based on results from individual FP strain measurements (Figure 5), as well as combinations previously used in literature. FRET pairs such as the combination of a cyan and yellow FP are often used in biosensors or other FRET-applications such as protein localization studies. In this study, the FRET-pairs sAphrodite/sCerulean, which is used for example in a biosensor [40], and mVenus/mCerulean, were chosen. Non-overlapping FP combinations such as BFP/GFP/RFP are widely used for biosensors, cell population discrimination within within co-cultures and other multiplexed measurements [30, 29]. In this study, the pairs eGFP/LSSmOrange and eGFP/LSSmApple were chosen as examples for FP combinations used in Matryoshka biosensors [28, 52], as well as three alternative combinations of eGFP with either mTagBFP2, mScarlet-I or mCherry, respectively, which could be used as alternative reference FPs for Matryoshka sensors. While eGFP is more commonly used, our study confirmed claims from previous *in vivo* studies that mNeonGreen is generally brighter [53], and therefore superior for cyanobacterial applications. Therefore, mNeonGreen was combined with mTagBFP2, mScarlet-I, or mCherry and also included in the dual-FP evaluation pipeline. Dual FP constructs were measured alongside their respective single FP controls and the EVC, using cytometer settings and detector configurations suitable for monitoring both fluorescent channels (Supplementary File S2).

To compare the dual FP-combinations, scatter plots for each of these groups of strains were generated, with the fluorescence intensity measured with either the optimal or second best channel for FP 1 on the x-axis and FP 2 on the y-axis (Figure 7). For the mVenus/mCerulean pair (Figure 7 A), both combinations of orders in the operon were tested. For some dual FP combinations, B525-40-A, the second best channel, was used instead of B510-20-A, the optimal channel, due to limitations in the detector configuration. Still, both green FPs can easily be identified and clearly separated from the autofluorescence using either channel.

**Figure 7:**
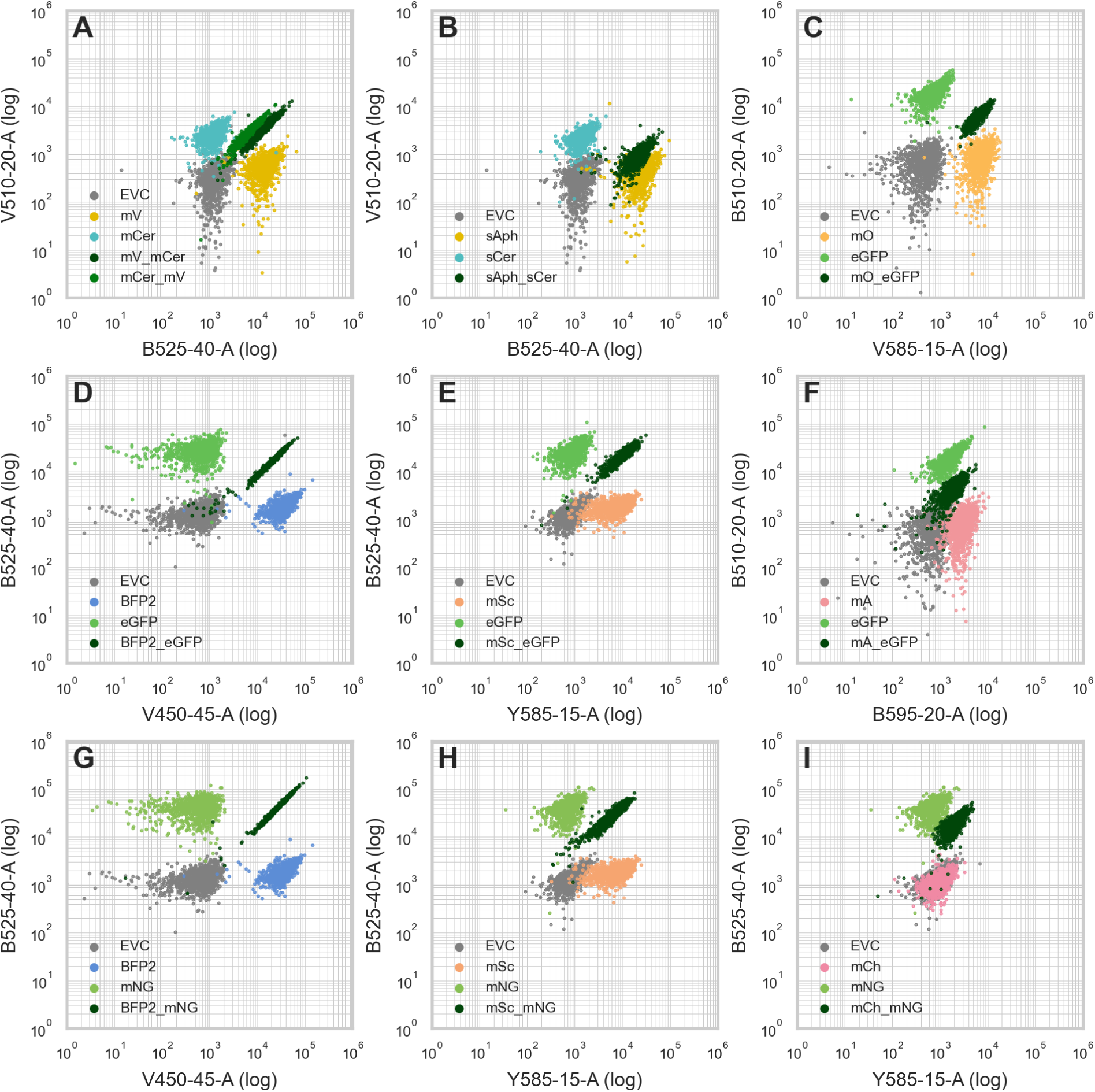
Dot plots of dual FP compared to respective single FP and EVC strains. A: mV (mVenus) and mCer (mCerulean), B: sAph (sAphrodite) and sCer (sCerulean), C: mO (LSSmOrange) and eGFP, D: BFP2 (mTag-BFP2) and eGFP, E: mSc (mScarlet-I) and eGFP, F: mA (LSSmApple) and eGFP, G: BFP2 (mTagBFP2) and mNG (mNeonGreen), H: mSc (mScarlet-I) and mNG (mNeonGreen), I: mCh (mCherry) and mNG (mNeonGreen). Populations of the EVC (empty vector control, in gray), single FP (FP-specific color) and dual FP strains (dark green) are shown representing 1,000 cyanobacterial singlet cells each. The logarithmic Y- and X-axes represent the optimal channel (or second best channel for eGFP and mNG due to restrictions in the detector configuration) for the first and second FP in the operon construct, respectively. In the channel names, the letters describe the excitation laser (V=Violet: 405 nm; B=Blue: 488 nm; Y=Yellow: 561 nm), -A stands for area of the signal peak, while numbers describe the emission wavelength and bandwidth of the filters. EVC: Empty vector control.

For the FRET pair mCerulean/mVenus, a clear separation of both single FP and dual FP strains from the autofluorescence of the EVC can be observed, with only a small overlap in the CFP channel V510-20-A. While both dual FP variants are mostly overlapping, mCerulean/mVenus exhibits a lower fluorescence signal in the B525-40-A channel than mVenus/mCerulean (Figure 7 A). The other FRET pair sAphrodite/sCerulean demonstrates similar results with the exception that the intensity of the sCerulean signal is slightly lower compared to the single FP sCerulean strain (Figure 7 B). The popular CFP/YFP FRET pair has already been successfully applied in form of an intramolecular biosensor to study cyanobacterial P_II_ [27].

Both BFP/GFP pairs, mTagBFP2/eGFP and mTagBFP2/mNeonGreen show very similar results with a clear separation of all cell populations and a very defined population of the dual-FP-strain (Figure 7 D, G). Similar observations apply for the GFP/RFP pairs, eGFP/mScarlet-I and mNeonGreen/mScarlet-I where only the mScarlet-I population is partly overlapping with the EVC (Figure 7 E, H). Still, this GFP/mScarlet pair is the only GFP/RFP pair that has already successfully been applied in a cyanobacterial study for tracking separate plasmids via microscopy [18].

The LSSmOrange/eGFP pair shows a clear separation of all four populations with only a small overlap of the dual-FP and LSSmOrange population (Figure 7 C). This overlap is caused by a slightly lower fluorescent intensity of the eGFP in the dual-FP-population which can also be found in the LSSmApple/eGFP and mCherry/mNeonGreen populations (Figure 7 F, I). As for the YFP/CFP pairs, the most likely explanation is a reduced expression or translation of the second FP in the operon. A FRET effect as a possible explanation for this phenomenon is less likely as eGFP and LSSmOrange/LSSmApple do not form a FRET pair and because for the mCerulean/mVenus combination the acceptor FP emission would be reduced (Figure 7 A, B, C, F).

Of the tested combinations, only the mCherry/mNeonGreen pair is not suitable, as the mCherry is overlapping almost completely with the EVC population (Figure 7 I). Other suitable FP combinations, that were only tested as single FP combinations in this study are for example YFP/BFP, BFP/RFP, and YFP/RFP combinations (Figure S8 A, B, C). Dual BFP/YFP systems have already been successfully applied in synthetic microbiology studies including cyanobacteria [16, 17]. Interestingly, also the combination of CFPs with mNeonGreen is potentially a good option (Figure S8 D), because the excitation spectrum of mNeonGreen is shifted a bit up compared to eGFP (Figure S1), for which a combination with a CFP is not possible due to co-excitation and detection in the V510-20-A channel (Figure S9 B). This combination of mTurquoise-tagged Rubisco has previously been used simultaneously with mNeonGreen-tagged components of proteins involved in its spatial distribution [24]. Combinations with LSS-FPs are usually a good option (e.g. YFP/LSSmOrange or BFP/LSSmApple, Figure S8 E, F) as long as they don’t have a spectral overlap in both the emission and excitation spectrum such as the combination BFP or CFP with LSSmOrange (Figure S9 D, E). Combinations of GFP with LSSmOrange or LSSmApple are for example used in Matryoshka biosensors [28, 52]. Combinations with spectral overlaps in emission and excitation spectra such as BFP/CFP, CFP/GFP or GFP/YFP combinations are generally unsuitable (Figure S9 A, B, C). Interestingly, the combination of a cyan FP such as mTuquoise2 with mScarlet-I is not suitable due to an increase signal of mScarlet-I in the V510-20-A channel (Figure S9 F), although neither the excitation nor the emission spectra of both FPs overlap much (Figure S1).

In general, FP combinations are most likely suitable if they do not have a spectral overlap in the excitation as well as the emission spectra, so they have a unique channel in the cytometer, and are strong enough to be distinguishable from the autofluorescence, showing a high signal-to-background ratio (Figure 5, Figure S6). If used in separate cells (e.g. in co-cultures) even combinations with big spectral overlaps in excitation and emission can be differentiated applying single-cell flow cytometry as long as they do not have the same optimal channel.

### Concluding assessment

In both the cytometry and plate reader measurements, green and yellow FPs are generally well distinguishable from the background (Figure 4, Figure 5). While they may remain the preferred choice when using a plate reader, blue FPs, especially mTagBFP2, perform even better under suitable cytometer conditions. With the exception of mCherry and mKate2, all FPs tested in this work are measurable using either plate reader or cytometry and, more importantly, distinguishable from the cyanobacterial background fluorescence, and are therefore suitable depending on the final application and measurement settings available.

Under optimized conditions and with the appropriate equipment, multiple dual FP combinations could be co-expressed and distinguished. Both eGFP and mNG can be distinctly observed together with either BFPs or LSS-FPs/RFPs (Figure 7). It should be noted that eGFP and mNG, while both green FPs, vary slightly in their emission maxima. This may be of use when choosing a green partner for BFP or RFP dual FP constructs. The same is true for mTagBFP2 and Electra2, although the latter was not tested in a dual combination in this work. It should be kept in mind that all FPs tested in this work were tested under a strong constitutive promoter. While some of them were also shown to work with much weaker or inducible promoters in other works [12, 31, 54], FPs should be carefully selected and tested against a control strain under these circumstances. Despite using the same genetic background and context, this is not necessarily a guarantee for equal expression and protein levels [19]. Therefore, ratios published in this work are subject to change and optimization, dependent on individual design and genetic context.

Overall, 12 out of the 14 experimentally tested FPs could be reliably distinguished from cyanobacterial autofluorescence using two distinct methods of measurement. Out of 10 dual FP combinations, 8 could be separated from their single FP counterparts. To the best of our knowledge, this is the first demonstration of the two LSS FPs, as well as the RFP mScarlet-I, in a cyanobacterium in a quantitative and comparative fashion. This study provides a valuable resource for future fluorescence-based cyanobacterial research.

## Acknowledgements

We would like to thank Daniel C. Ducat for gifting us mNeonGreen, St. Elmo Wilken for gifting us Electra2 and mScarlet-I, Joana C. Pohlentz (AG Feldbrügge, Institute for Microbiology, HHU) for gifting us mKate2, the Center for Advanced Imaging (CAi) and Center for Structural Studies (CSS) at HHU as well as CRC1535 MibiNet project Z01 for gifting us mCherry, and Mayuri Sadoine for gifting us eGFP, sAphrodite, sCerulean, LSSmApple, LSSmOrange and mTurquoise2. We highly appreciate the help of Dennis Dienst and St. Elmo Wilken with proofreading and valuable feedback.

## Author contributions

D.H., I.M.A., and A.B. conceptualized and supervised the project. D.H. and A.B. formally analyzed the data and wrote the original draft. D.H., J.B., A.G., and F.S. performed experiments and curated the data. D.H., I.M.A. and A.B. reviewed and edited the final draft. I.M.A acquired the funding and was responsible for project administration. All authors contributed to revision of the draft and gave their final approval.

## Declaration of generative AI and AI-assisted technologies in the writing process

The authors would like to explicitly state that no generative AI was used in the writing process. GitHub Co-Pilot was used in Visual Studio Code for help with implementation of the code for data processing and visualization. All outputs were reviewed and confirmed carefully.

## Data Availability

All raw data files, including .fsc files, were deposited in an Annotated Research Context (ARC) and will be made available once peer-reviewed. DNA coding- and amino acid sequences of all FPs used in this work are attached as fasta files (Supplementary Files 3 and 4) as well as full plasmid maps, which are attached as .gb files in Supplementary file 5.

## Supplementary Information for

### Overview of Supplementary Files in this Work

- Supplementary File 1 (.pdf): Supplementary Methods and Figures
- Supplementary File 2 (.xlsx): Plate Reader and Flow Cytometry settings used in this work.
- Supplementary File 3 (.fa): Fasta files of all FP DNA sequences used in this work.
- Supplementary File 4 (.fa): Fasta files of all FP amino acid sequences used in this work.
- Supplementary File 5 (.zip): Genbank files of all plasmids constructed in this work.

### Overview of Supplementary and Figures

- Supplementary Figure S1: Excitation and emission spectra of all FPs tested in this work, as deposited on FPbase.org. (FPbase IDs can be found in the main text, Table 1)
- Supplementary Figure S2: Excitation and emission scans of the wild type *Synechocystis* strain.
- Supplementary Figure S3: Emission scans of sCerulean and mKate2 strains.
- Supplementary Figure S4: OD_750_- and blank-corrected emission scans of each FP strain and corresponding EVC at optimal settings.
- Supplementary Figure S5: OD_750_-corrected fluorescence intensity of each FP strain and EVC at optimal settings.
- Supplementary Figure S6: Signal-to-background ratio_cyt_ for dual FPs.
- Supplementary Figure S7: Split violin plots of single RFP populations compared to the EVC in the four suitable bandpass filters.
- Supplementary Figure S8: Dot plots of other suitable dual FP combinations visualized by the respective single FP and EVC strains.
- Supplementary Figure S9: Dot plots of other unsuitable dual FP combinations visualized by the respective single FP and EVC strains.

**Figure S1:**
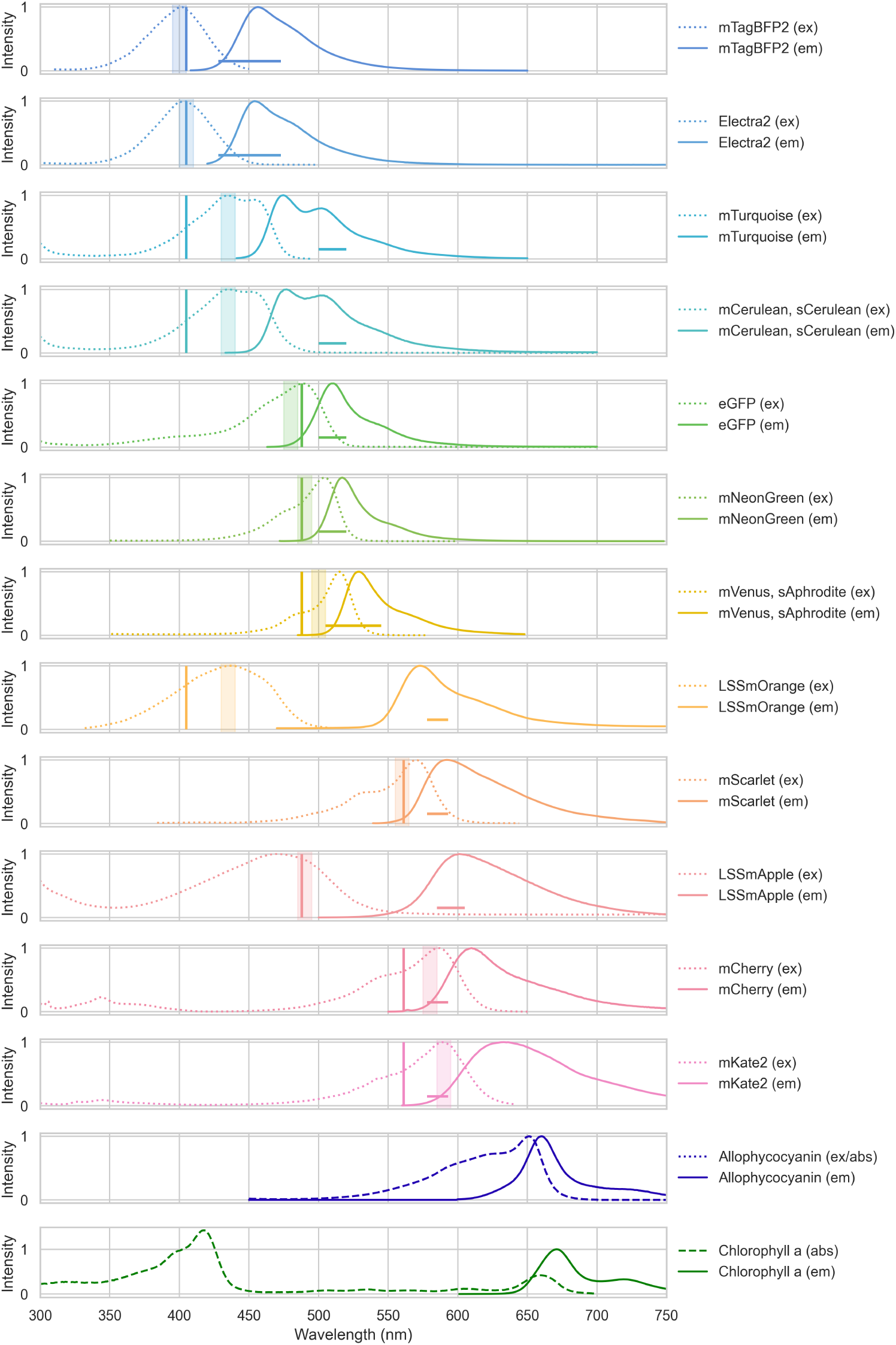
Excitation and emission spectra of all FPs tested in this work, as found on FPbase.org. (FPbase IDs can be found in the main text, Table 1) Dotted lines show excitation, continuous lines emission spectra. Excitation ranges used in plate reader measurements throughout this work are shadowed. Vertical lines represent excitation lasers used in cytometer measurements, horizontal lines represent the range of optimal bandwidth filters used in cytometer measurements. Allophycocyanin and Chlorophyll a excitation, emission, and absorption spectra are shown at the bottom. Note: The excitation and absorption spectra of allophycocyanin overlap exactly.

**Figure S2:**
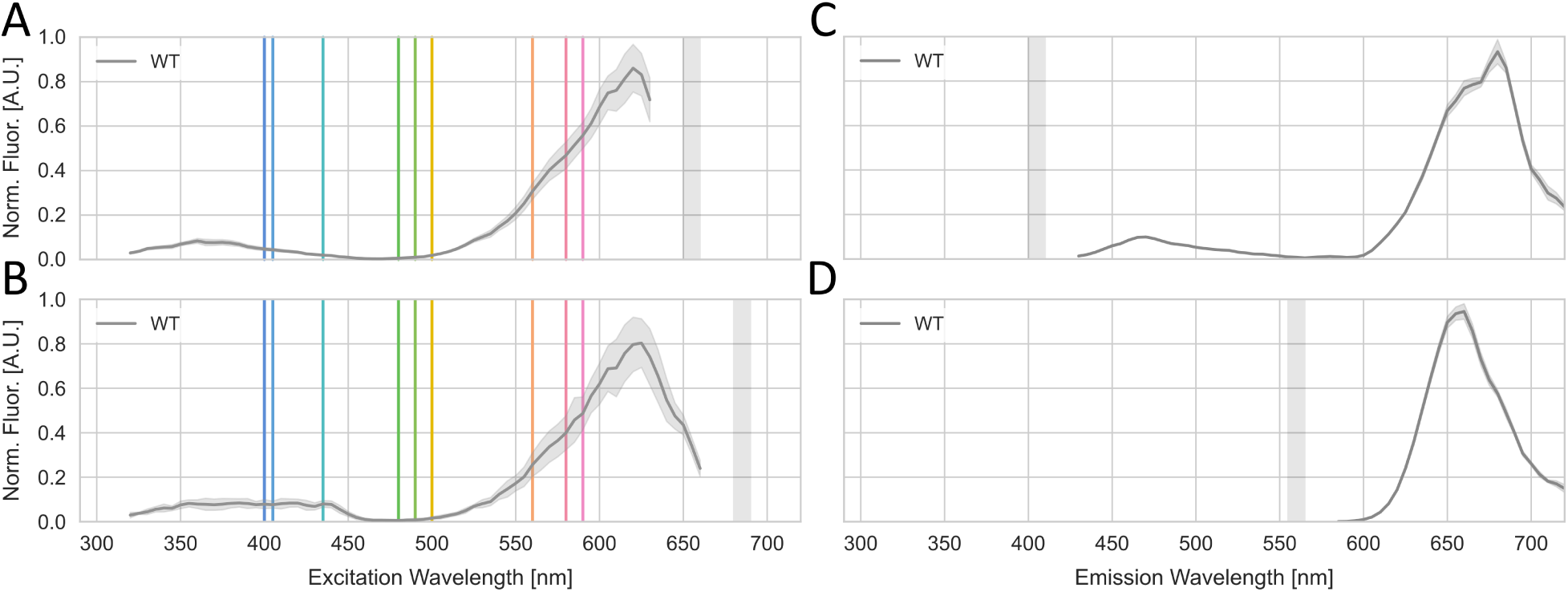
Excitation and emission scans of the wild type *Synechocystis* strain. WT: Wild type *Synechocystis*. Norm. Fluor. [A.U.]: Normalized Fluorescence [arbitrary units]. A,B: Whole cell excitation scans, measuring emission at either A. 655 nm (+/- 5 nm), or B. at 685 nm (+/- 5 nm). The corresponding emission wavelength is shaded in grey. C,D: Whole cell emission scans using excitation wavelength at either C. 405 nm (+/- 5 nm), or D. at 560 nm (+/- 5 nm). Raw data were normalized to OD_750_. Data for each plot was scaled ranging from 0 to 1. The shaded region shows the standard deviation of four analyzed replicates. Colored vertical lines represent excitation wavelengths used for different FPs throughout this work.

**Figure S3:**
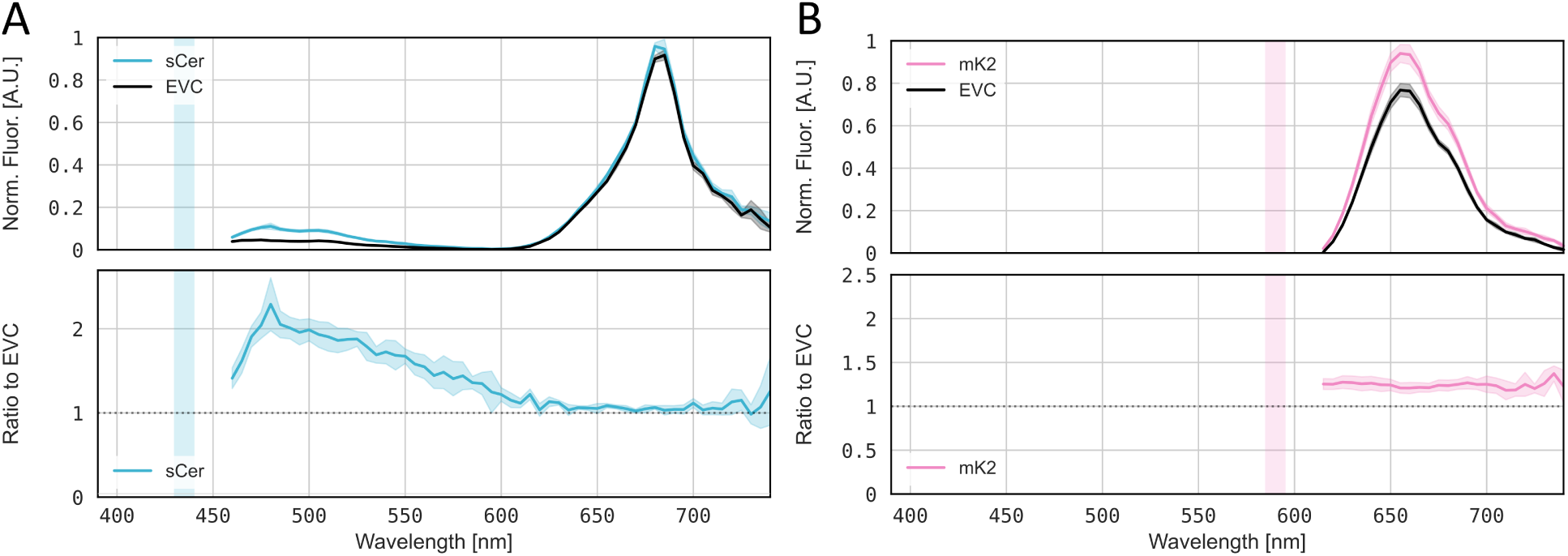
Emission scans of sCerulean and mKate2 strains. A. sCer: sCerulean, B. mK2: mKate2. A,B, top panels: Whole cell emission scan of *Synechocystis* expressing FP of interest. For each condition, the empty vector control (EVC) strain was measured alongside. Raw data were normalized to OD_750_. Data for each plot (FP+ respective EVC in black) was scaled ranging from 0 to 1, The shaded region shows the standard deviation of four analyzed replicates. A,B, bottom panels: Ratio of FP-encoding strain to EVC; the shaded region shows the standard deviation of four analyzed replicates. The optimal range of excitation wavelengths used for each FP according to published data is marked by a vertical shaded bar.

**Figure S4:**
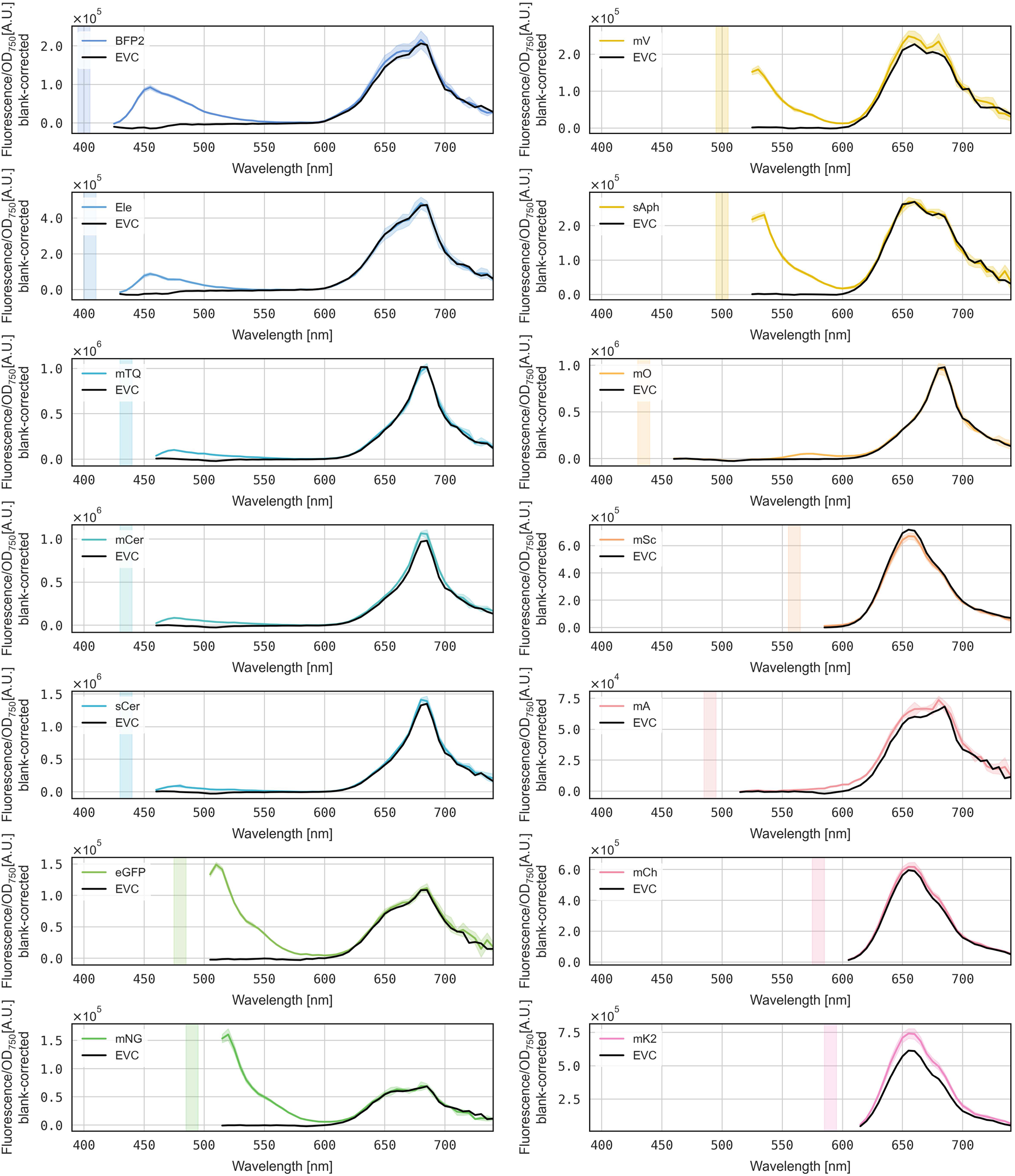
Blank-corrected emission scans of all single-FP strains. Whole cell emission scan of *Synechocystis* expressing FP of interest. For each condition, the empty vector control (EVC) strain was measured alongside. Data for each plot was blank-corrected using BG11 medium, then normalized to OD_750_. For each strain, 4 replicates were analyzed. BFP2: mTagBFP2. Ele: Electra2. mTQ: mTurquoise2. mCer: mCerulean. sCer: sCerulean. mNG: mNeonGreen. mV: mVenus. sAph: sAphrodite. mO: LSSmOrange. mSC: mScarlet-I. mA: LSSmApple. mCh: mCherry. mK2: mKate2.

**Figure S5:**
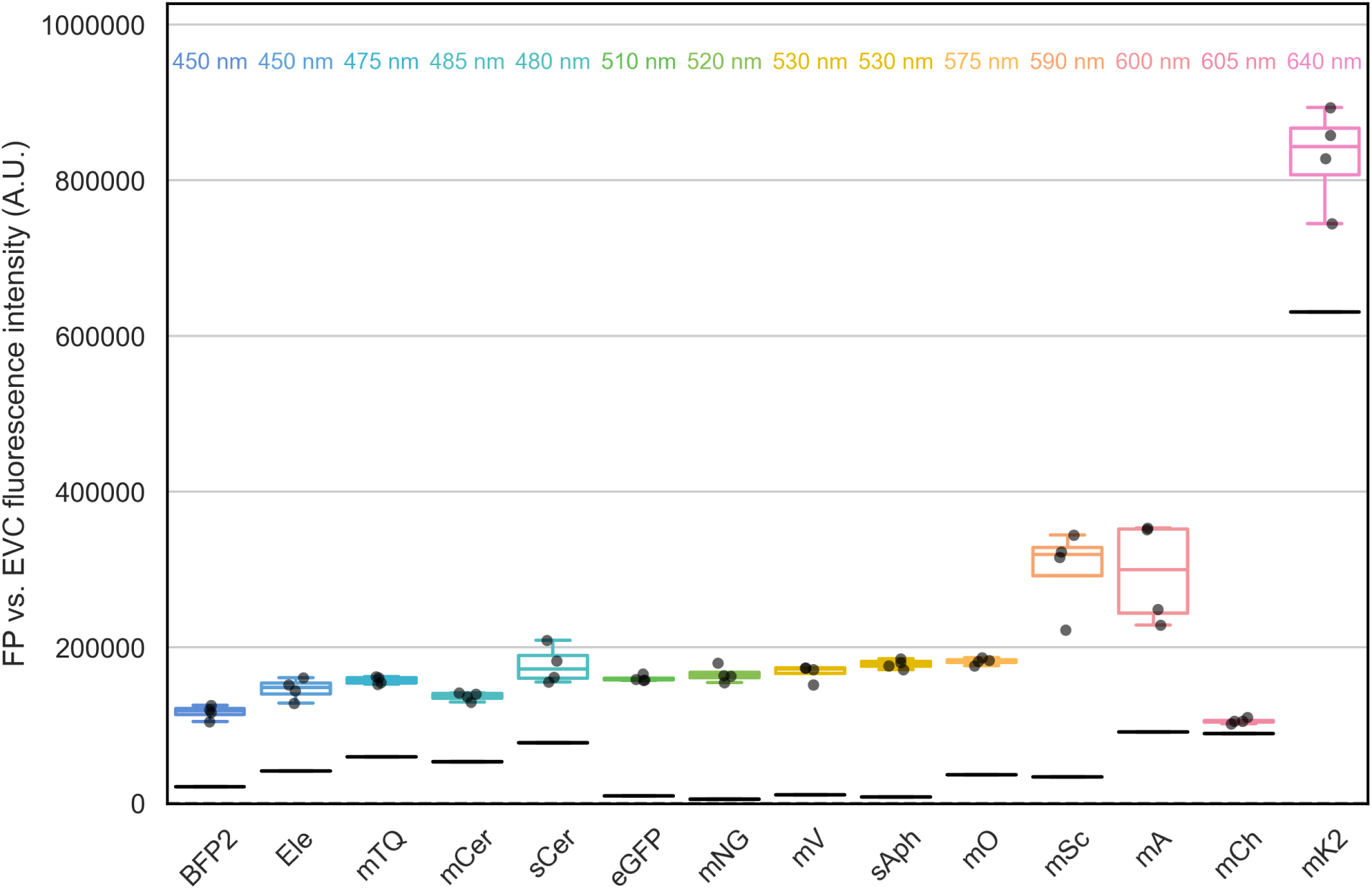
**OD**_750_**-corrected fluorescence intensity of each FP strain and EVC at optimal settings.** Fluorescence data used for determination of optimal signal-to-background ratios_PR_, corresponding to Figure 4. Using the optimum emission wavelength, (shown on top, +/- 5 nm), fluorescence intensity was determined for each FP and the empty vector control (EVC, black). Data represents four biological replicates. EVC values are different between samples due to different gain settings and autofluorescence. BFP2: mTagBFP2. Ele: Electra2. mTQ: mTurquoise2. mCer: mCerulean. sCer: sCerulean. mNG: mNeonGreen. mV: mVenus. sAph: sAphrodite. mO: LSSmOrange. mSC: mScarlet-I. mA: LSSmApple. mCh: mCherry. mK2: mKate2.

**Figure S6:**
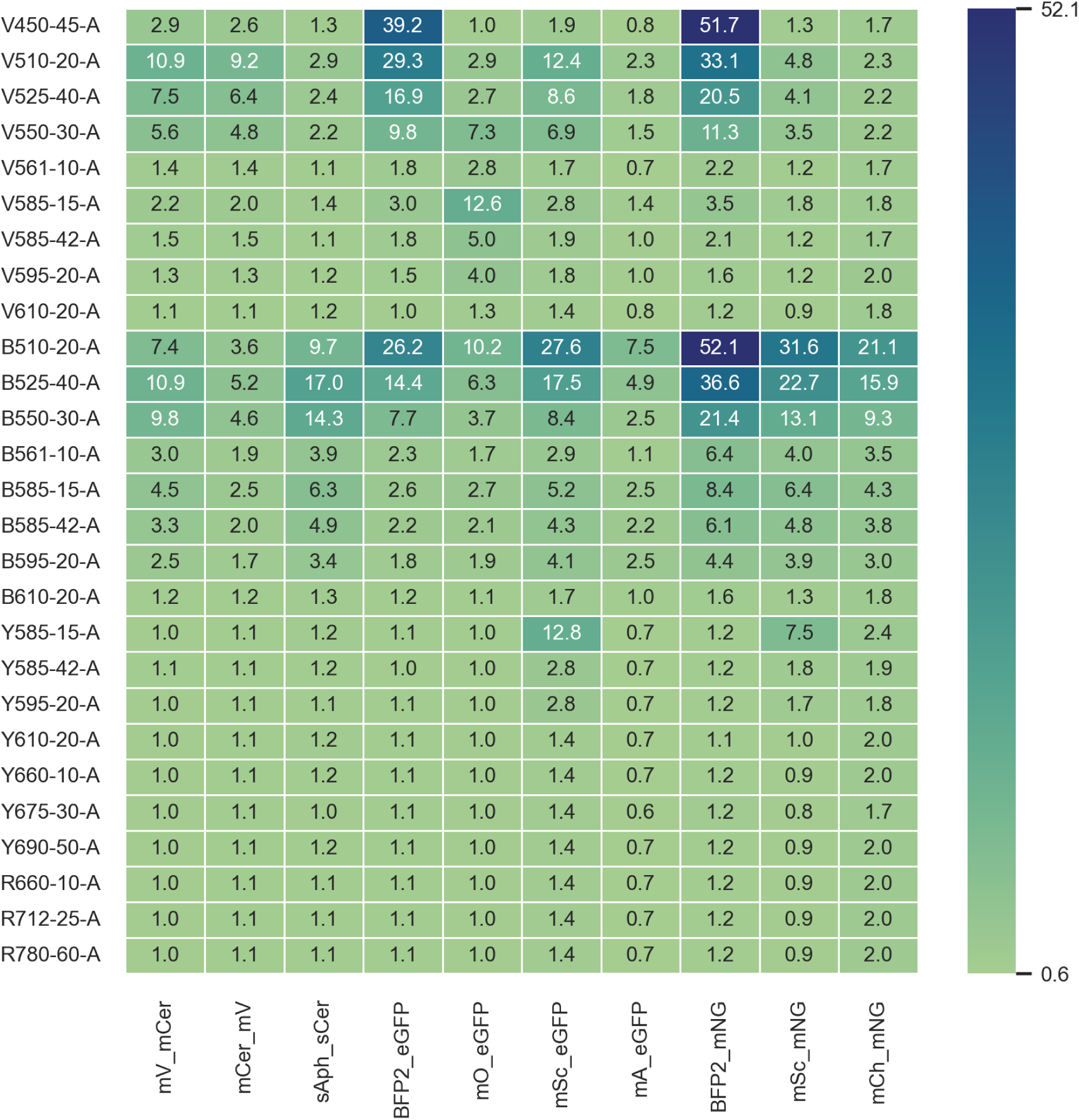
Signal-to-background ratio_cytometer_ for dual FPs. The ratio of median FP/EVC (empty vector control) fluorescence (signal-to-background ratio_cyt_) was determined for all combinations of dual FP-strains (x-axis) and available fluorescence channels (y-axis). In the channel names, the letters describe the excitation laser (V=Violet: 405 nm; B=Blue: 488 nm; Y=Yellow: 561 nm; R=Red: 638 nm), -A stand for area of the signal peak, while numbers describe the emission wavelength and bandwidth of the filters. Color scale represents a 0.6 - 52.1 scale of the signal-to-background ratio_cyt_. BFP2: mTagBFP2. mCer: mCerulean. sCer: sCerulean. mNG: mNeonGreen. mV: mVenus. sAph: sAphrodite. mO: LSSmOrange. mSC: mScarlet-I. mA: LSSmApple. mCh: mCherry.

**Figure S7:**
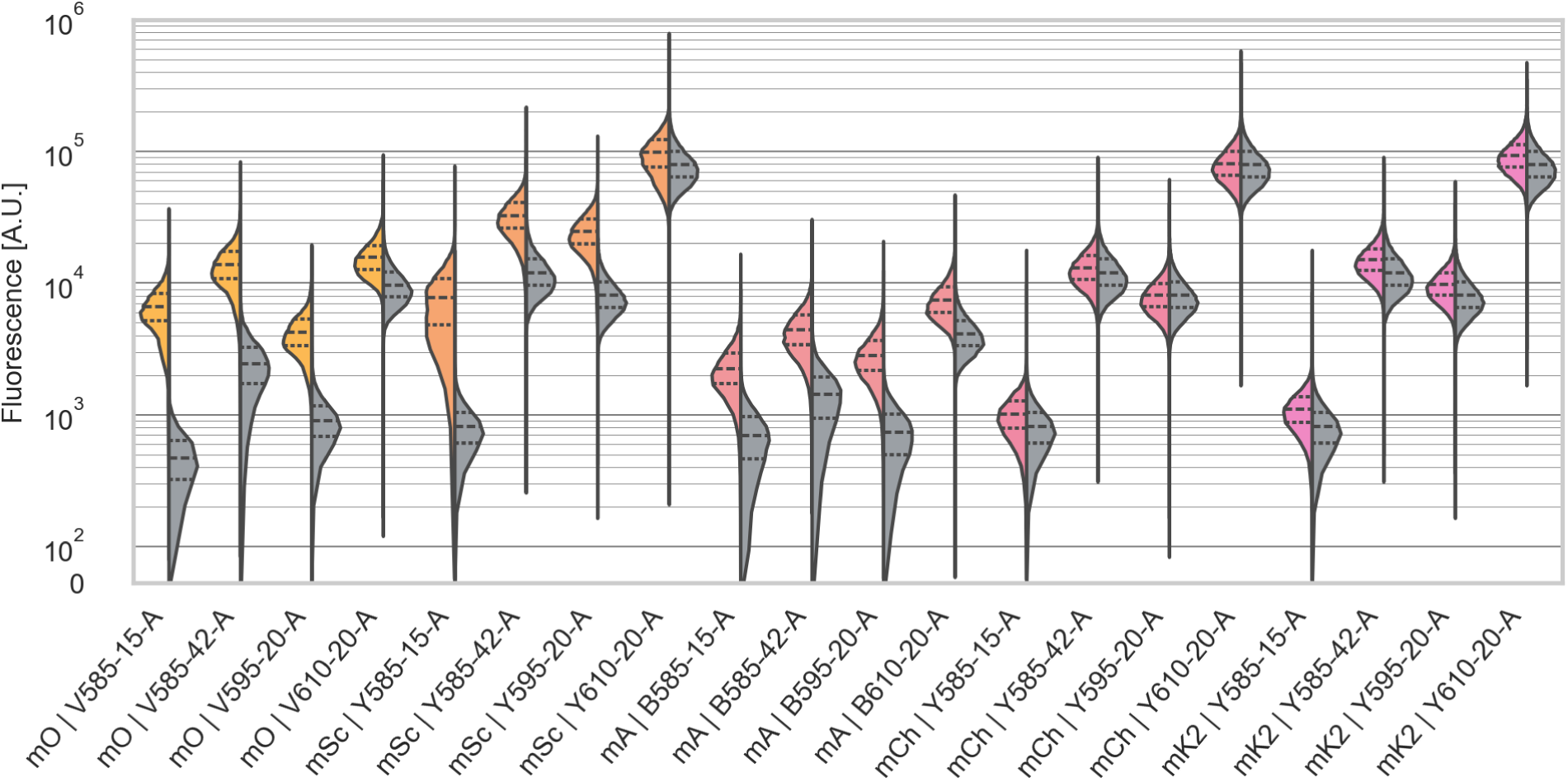
Split violin plots of single RFP populations compared to the EVC in the four suitable bandpass filters. Each pair was measured with one of the four suitable bandpass filters for RFPs in their respective excitation laser. Violin plots show the kernel density estimation (KDE) and quartiles calculated on all cyanobacterial singlet cells (max. 50,000) for each FP population (left side of violin) and the corresponding EVC (right side of violin). Dashed lines represent the median (50th percentile) and dotted lines represent the 25th and 75th percentile. Violin plots are displayed on a symlog scale with a linear region from 0 - 10^2^ and a logarithmic region from 10^2^ - 10^6^ showing the fluorescence [arbitrary units]. In the channel names, the letters describe the excitation laser (V=Violet: 405 nm; B=Blue: 488 nm; Y=Yellow: 561 nm), -A stand for area of the signal peak, while numbers describe the emission wavelength and bandwidth of the filters. EVC: Empty vector control. mO: LSSmOrange. mSC: mScarlet-I. mA: LSSmApple. mCh: mCherry. mK2: mKate2.

**Figure S8:**
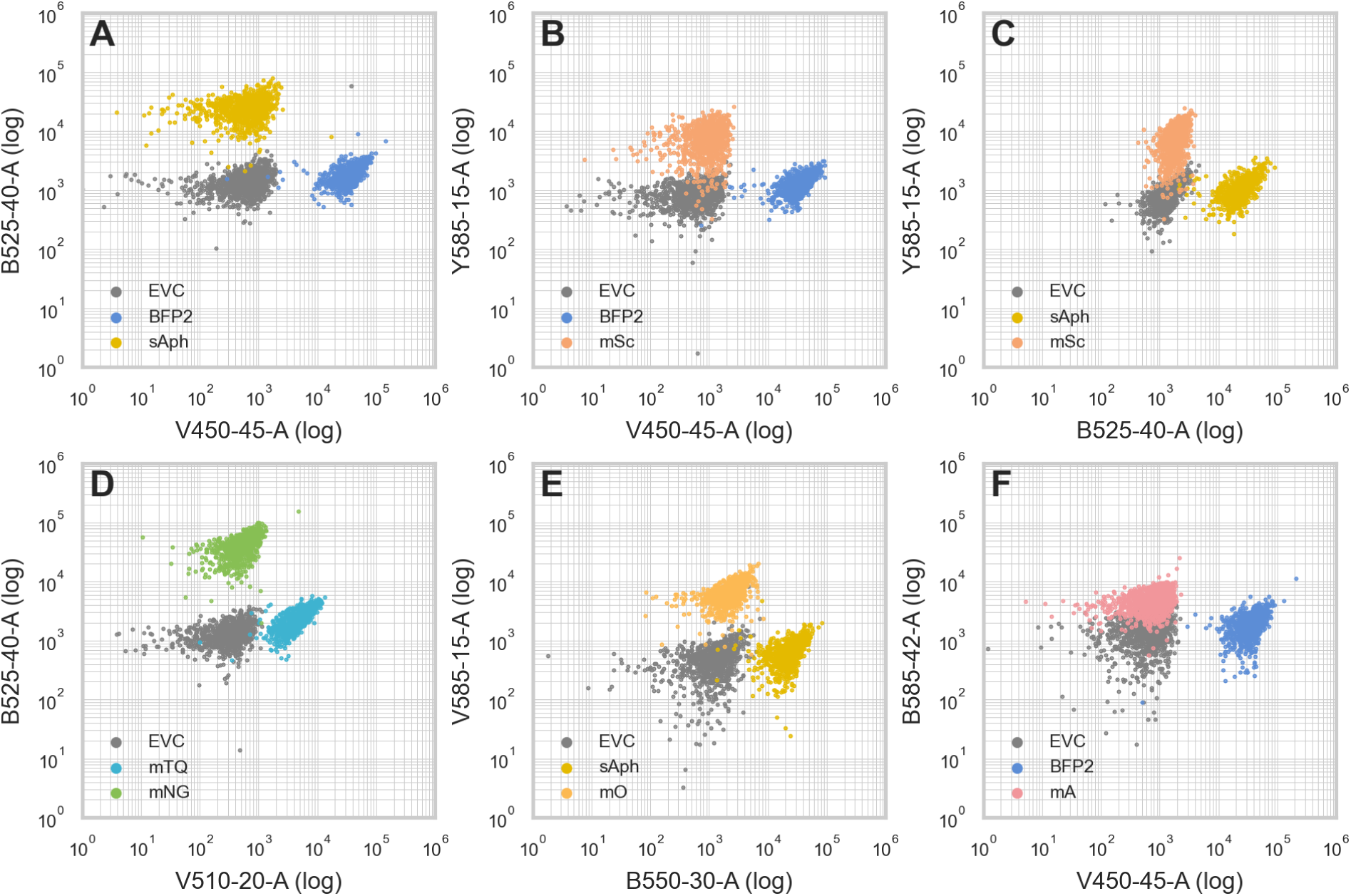
Dot plots of other suitable dual FP combinations visualized by the respective single FP and EVC strains. A: BFP2 (mTagBFP2) and sAph (sAphrodite), B: BFP2 (mTagBFP2) and mSc (mScarlet-I), C: sAph (sAphrodite) and mSc (mScarlet-I), D: mTQ (mTurquoise2) and mNG (mNeonGreen), E: sAph (sAphrodite) and mO (LSSmOrange), F: BFP2 (mTagBFP2) and mA (LSSmApple). Populations of the EVC (empty vector control, in gray), single FP (FP-specific color) are shown representing 1,000 cyanobacterial singlet cells each. The logarithmic Y- and X-axes represent the optimal channel (or second best channel for mNG, sAph or mA due to restrictions in the detector configuration) for both FPs. In the channel names, the letters describe the excitation laser (V=Violet: 405 nm; B=Blue: 488 nm; Y=Yellow: 561 nm), -A stands for area of the signal peak, while numbers describe the emission wavelength and bandwidth of the filters.

**Figure S9:**
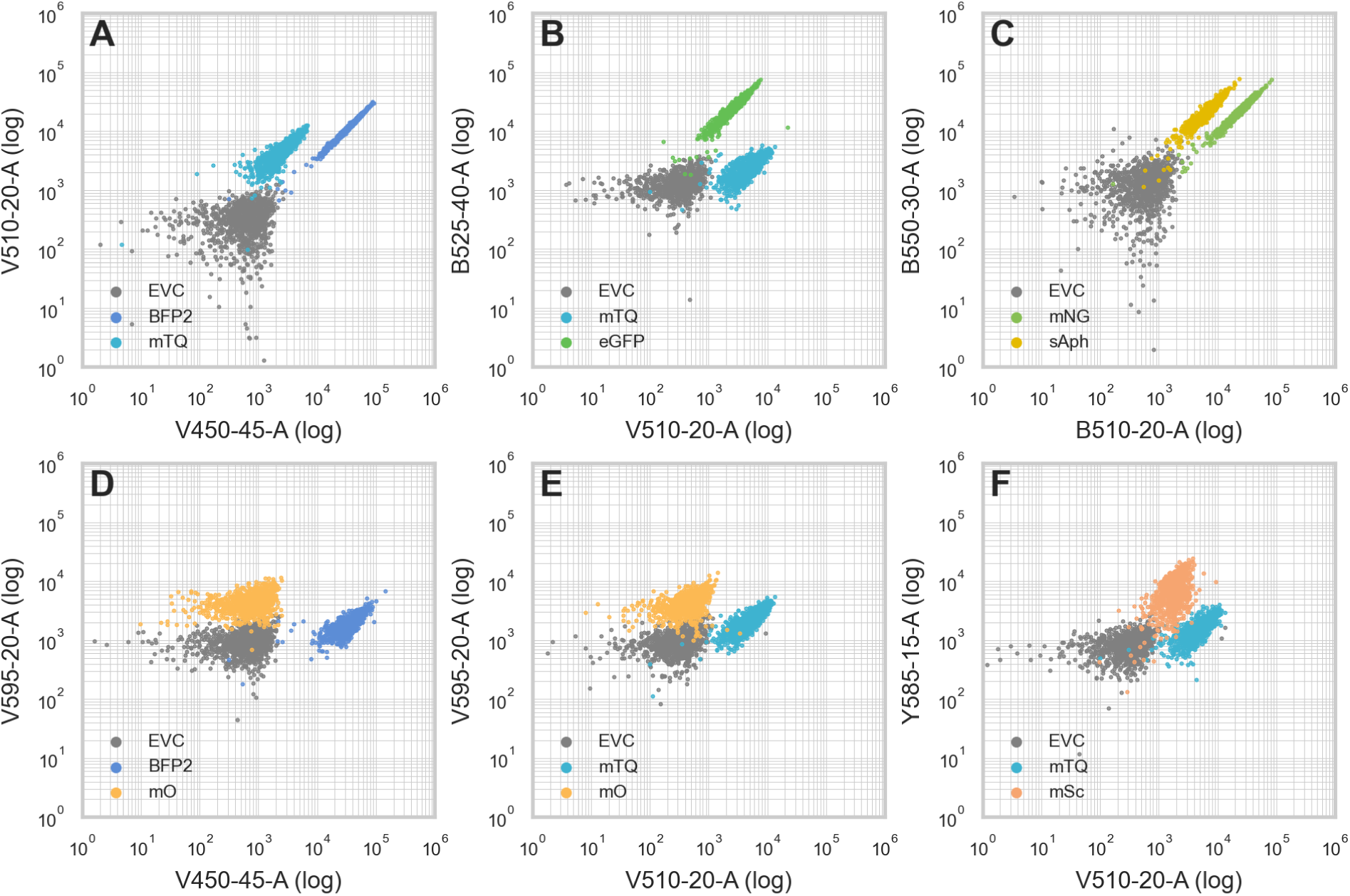
Dot plots of other unsuitable dual FP combinations visualized by the respective single FP and EVC strains. A: BFP2 (mTagBFP2) and mTQ (mTurquoise2), B: mTQ (mTurquoise2) and eGFP, C: mNG (mNeonGreen) and sAph (sAphrodite), D: BFP2 (mTagBFP2) and mO (LSSmOrange), E: mTQ (mTurquoise2) and mO (LSSmOrange), F: mTQ (mTurquoise2) and mSc (mScarlet-I). Populations of the EVC (empty vector control, in gray), single FP (FP-specific color) are shown representing 1,000 cyanobacterial singlet cells each. The logarithmic Y- and X-axes represent the optimal channel (or second best channel for eGFP, sAph, and mO due to restrictions in the detector configuration) for both FPs. In the channel names, the letters describe the excitation laser (V=Violet: 405 nm; B=Blue: 488 nm; Y=Yellow: 561 nm), -A stands for area of the signal peak, while numbers describe the emission wavelength and bandwidth of the filters.

## Supplementary methods

### Plate Reader measurements

Samples of 200 µL cyanobacterial culture were loaded in a black-walled, clear-bottom 96-well plate. Before each measurement, plates were shaken for 10 seconds at 500 rpm (double orbital). Excitation and emission scans of all samples were performed in 5 nm increments with separate settings for each strain. In addition, absorbance scans between 220 - 1000 nm, including the absorbance at 750 nm was recorded for all wells. For each FP, the optimal excitation wavelength, +/- 5 nm, was selected based on published values or the excitation maximum (Figure S1). In some cases, for example eGFP, mNG, and mVenus/sAphrodite, it was necessary to select an excitation range outside of the optimum to prevent an overlap between excitation and emission of the FP. Based on these excitation settings, an individual gain- and focal height adjustment using the emission maximum was performed separately for each FP and stored as a separate protocol. The gain was set to a target value of 80 % of the maximum dynamic range of the plate reader. In some cases, the optimal gain adjustment for an FP led to overflow values in the autofluorescence range. In these cases (excitation wavelengths 560, 580, 590 nm), an additional protocol was created with gains set to an emission wavelength of 650 nm, resulting in overflow-free values across the entire spectrum. These were also stored as separate protocols. All measurement protocols were run across the entire range of FP-strains, recording data for all FP/protocol combinations in four replicates each. The EVC and WT controls were included in each case. For Figures 1 - 3 and S2, data from either FP-optimized or overshoot-free (LSSmOrange from mCerulean-optimized protocol, LSSmApple from mNeonGreen-optimized protocol, mCherry, mKate2, mScarlet-I from respective autofluorescence-optimized protocols) are plotted and shown. For calculating the optimal signal-to-background ratio_PR_ in Figure 4, data from each optimal protocol was used, even if overshoots were recorded. The reason for this is that in practical applications, the settings would be optimized to gain the best measurements within the FP-relevant wavelengths, not across the entire spectrum. All data sets, including those not shown in any Figures, were deposited. The settings for the excitation scans shown in Figure S2 were selected in the same manner. All protocols and their settings are summarized in Supplementary File S2, tab 1.

